# OCTRON: a general purpose segmentation and tracking pipeline for behavioral experiments

**DOI:** 10.64898/2025.12.20.695663

**Authors:** Ragnhild Irene Jacobsen, Nadia M. van Eekelen, Mikkel Roald-Arbøl, Sofía Miñano, Laurel Humphrey, Johnston Renton, Takato Honda, Elke van Rooij, Jason Rivera, Oscar M. Arenas, Ellen A. Lumpkin, Sofia Maccuro, Kendra C. Buresch, Eve Seuntjens, Horst A. Obenhaus

## Abstract

Quantifying animal behavior across species with complex, deformable body plans requires capturing full-body shape rather than sparse skeletal keypoints. Yet accessible, end-to-end tools that take researchers from mask annotation through model training to reliable multi-object tracking are lacking. Here we present OCTRON, a fully trainable pipeline for multi-object segmentation, detection, and tracking that is broadly applicable across experimental setups and organisms. OCTRON leverages large segmentation foundation models to rapidly generate experimenter-supervised annotations from minimal input, overcoming the manual labeling bottleneck of supervised segmentation. These annotations train compact, task-specific models: segmentation networks for extracting pixel-level information such as shape and appearance, or lightweight detection networks for fast centroid tracking. Coupled with multi-object trackers and an interactive track-curation interface in napari, OCTRON enables complete behavioral quantification with minimal manual intervention. We validate OCTRON across species from transparent marine annelids to camouflaging cuttlefish, and in multi-species behavioral assays.

## Main

Automated video tracking has transformed the quantitative study of animal behavior, largely through deep-learning tools that infer position and pose from sparse anatomical keypoints. While frameworks such as DeepLabCut^1^, SLEAP^2^, and LightningPose^3^ achieve excellent performance in vertebrates and arthropods, their reliance on fixed skeletal landmarks makes them poorly suited for organisms with highly deformable body plans, such as hydra or cephalopods, that lack rigid joints or invariant reference points. In contrast, instance segmentation captures the full outline and spatial extent of anatomical regions, providing a much richer representation than sparse key points when body geometry itself changes continuously. Beyond quantifying full-body deformations, segmentation generates pixel-accurate masks of tracked body regions that can be applied directly to the raw video to extract appearance features such as color, luminance, and texture - unlocking levels of behavioral detail inaccessible to coordinate-based tracking.

However, generating segmentation data is extremely time-consuming, requiring users to manually delineate exact object boundaries frame by frame - a major bottleneck limiting adoption of segmentation-based tracking. To address this, we built OCTRON, a modular pipeline leveraging large segmentation foundation models to dramatically reduce the annotation burden. Specifically, we relied on SAM models (Segment Anything Model 2 and 3^4–6^). These models can rapidly generate high-quality masks from minimal user input, creating large training datasets in minutes. These curated annotations can then be used to train compact, domain-specific models - either segmentation models for detailed feature extraction or detection-only models (You Only Look Once [YOLO] model families 11 and 26^7,8^, Real-Time DEtection TRansformer [RT-DETR]^9^) for fast tracking optimized to the experimental context. Using these lighter, task-specific models, as opposed to the foundation models themselves, enables both high-resolution behavioral analysis and real-time processing of high frame-rate video.

Once frame-by-frame predictions have been made, maintaining coherent identity assignments across frames becomes critical, especially in multi-animal experiments as animals move, interact, and occlude one another. OCTRON integrates state-of-the-art multi-object trackers (BoxMOT^10^) that associate detections across frames using appearance and motion cues. BoxMOT’s flexible framework supports multiple tracking algorithms (ByteTrack^11^, BoT-SORT^12^, OC-SORT^13^, Hybrid-SORT^14^, Deep OC-SORT^15^, BoostTrack^16^), enabling users to select trackers optimized for their specific experimental needs, from single-subject analysis to crowded multi-animal scenarios. In addition, since trackers are prone to split up tracks when crossings or occlusions occur, OCTRON provides an interactive, user-friendly prediction cleaner widget that allows users to inspect and join track fragments.

At each stage - annotation, training, and tracking - OCTRON offers multiple models as plug-and-play solutions, ensuring broad applicability across diverse tracking scenarios. OCTRON also caters to the different levels of programming expertise that researchers have. By packaging state-of-the-art tools into an intuitive and familiar napari-based^17^ interface, OCTRON brings cutting-edge segmentation-based tracking within reach of researchers without a computational background, while a complete command-line interface enables computation workflows for more experienced users. Comprehensive documentation and best practices are available at https://github.com/OCTRON-tracking and https://octron-tracking.github.io/OCTRON-docs.

## Results

### OCTRON tracks species with deformable, complex bodies

To demonstrate the versatility of the species-agnostic OCTRON pipeline (Fig. 1a, Fig. S1), we used it to segment and track a number of non-model organisms whose behavior is challenging to analyze with keypoint-based methods. These organisms include camouflaging cuttlefish, octopus paralarvae, transparent marine annelids and ctenophores, as well as highly morphable hydra (Fig. 1b,c).

**Fig. 1:**
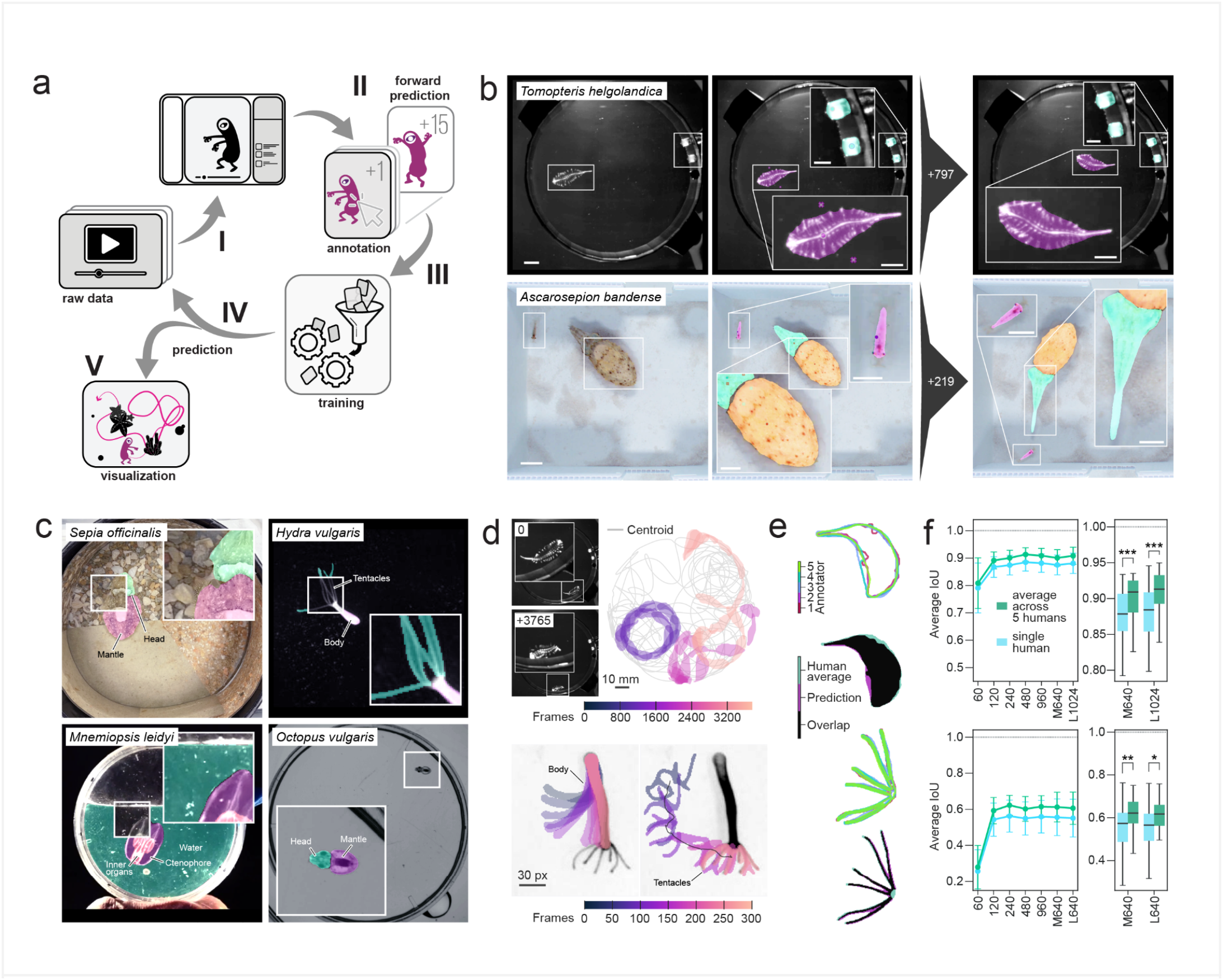
OCTRON workflow and performance. **a,** Pipeline schematic: I) load raw data in napari; II) create labels and use forward prediction for rapid training data generation; III) train and evaluate a new model; IV) predict on novel data with optional refinement; V) visualize results in OCTRON and/or use OCTRON library to work with the output. **b,** Annotation workflow examples. Top: *Tomopteris helgolandica* (pink) and LEDs (green). Scale bars: 10 mm and 5 mm (insets). Bottom: *Ascarosepion bandense* (green and orange) and shrimp (*Palaemonetes spp.,* pink). Scale bars: 25 mm and 12 mm (insets). Round dots: masked object, crosses: background. Right: Unsupervised forward prediction yields stable predictions after annotation of only a few frames and thereby accelerates training data generation (+797 frames after last annotated frame for *T. helgolandica*, +219 frames for *A. bandense*). **c,** Additional annotation examples across species: *Sepia officinalis, Hydra vulgaris, Mnemiopsis leidyi, Octopus vulgaris* paralarva. **d,** Prediction examples: stable segmentation and centroid tracking of *T. helgolandica* (top) and *H. vulgaris* (bottom). *T. helgolandica* masks are shown across three distinct episodes, color-coded by time. Row order is maintained across panels (e) and (f). **e,** Outlines of manually drawn masks by five different human annotators (first and third rows, color-coded by annotator) and masks used for benchmarking mask precision (second and fourth rows; human average - green, OCTRON prediction - purple, overlap - black). **f,** Segmentation accuracy vs. human labels. Left: Mean ± SD of the Intersection over Union (IoU, i.e. overlap) between human annotations and OCTRON-predicted segments provided by models trained on varying training dataset sizes (YOLO M model trained on 60, 120, 240, 480, 960 frames) and on the full training dataset (YOLO M, and YOLO L at either 640px or 1024px max training image size). Right: box plots showing the IoU for two models trained on the full training dataset. Top row (*T. helgolandica*), model: M640, t-stat: −3.77, p-value: 0.0005***, model: L1024, t-stat: −3.874, p-value: 0.0004***. Bottom row *(Hydra vulgaris* - tentacle region), model: M640, t-stat: −2.998, p-value: 0.0046**, model: L640, t-stat: −2.684, p-value: 0.011*).

For each species, we first loaded the relevant videos into OCTRON. Next, we created labels for the objects to be tracked and selected these objects in one or a few frames with single clicks or user-drawn outlines (Video S1). This rapidly created a video-specific memory in SAM that was used to predict the selected mask on subsequent frames in that same video (Fig. 1b). With this forward batch prediction, we annotated hundreds of frames within minutes to yield large training datasets, with the possibility to refine masks if necessary. The generated training datasets were then used to train YOLO segmentation models (YOLO11m or YOLO11l). By applying the trained models on held-out data, we found that OCTRON was able to reliably segment and track individuals of all the six species considered (*Ascarosepion bandense, Hydra vulgaris, Mnemiopsis leidyi, Octopus vulgaris, Sepia officinalis,* and *Tomopteris helgolandica*; two are shown in Fig. 1d). We then compared OCTRON-predicted masks with those created manually by humans (ground truth annotations). When evaluated across increasing training set sizes, OCTRON-trained models reached near-peak accuracy with as few as 120-240 annotated frames and agreed more closely with the five-annotator consensus than with any individual annotator (Fig. 1e,f and Fig. S2), outperforming single human annotators.

### Detection and segmentation in OCTRON unlock multi-object tracking, shape information, and keypoint extraction

To demonstrate OCTRON’s performance in challenging multi-animal experiments, we segmented and independently tracked the heads and mantles of three octopus paralarvae (*Octopus vulgaris*) swimming in the same arena (Fig. 2a). Even without manual track curation or any skeletal constraint linking body parts, tracking remained consistent across repeated path crossings (Video S2): over full recordings, each mantle trajectory paired uniquely with a single head trajectory, showing near-zero correlation with the head trajectories of other individuals (Fig. 2b,c). Beyond tracking position, the predicted masks capture continuous changes in body shape and size. Subjecting aligned head and mantle masks of swimming octopus paralarvae to principal component analysis (PCA) revealed low-dimensional shape modes corresponding to distinct postures and behavioral states (Fig. 2d). While PC1 captured changes in the animals’ pitch relative to the camera, PC2 separated a compact body configuration from the streamlined, elongated form adopted during forward jet-propelled swimming bouts. The latter is indicative of a streamlined configuration during swimming bouts when the animals propel themselves forward. Indeed, a correlation with speed confirmed that the elongated body form (low PC2 score) corresponded to higher swimming speeds (Fig. 2e). Combined, we show that access to segmentation data unlocks two complementary strengths: robust multi-object tracking for reliable identity preservation and rich shape information for high-resolution behavioral and morphological analyses.

**Fig. 2:**
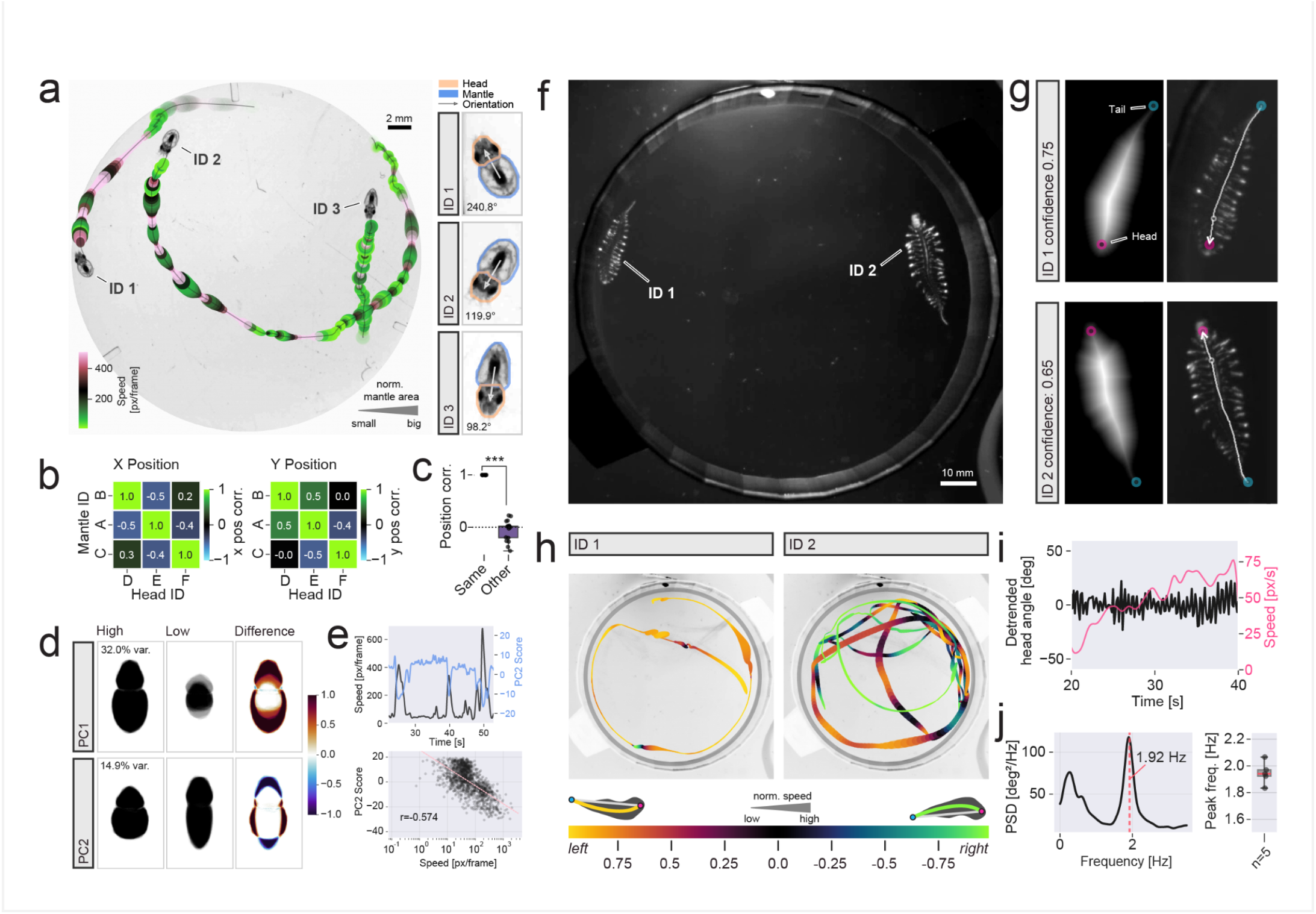
Multi-animal tracking and keypoint extraction. **a,** Octopus paralarvae open-field trajectories with size-coded mantle area and color-coded speed for three individuals. Right column: Zoomed in images of the same individuals, the arrow connects the centroid of the mantle (blue) to the head (orange). Heading angle is denoted in degrees in the bottom left. **b,** Position-correlation heatmaps (x, y) between mantle and head tracks; high-correlation pairs are denoted on the diagonal. **c,** Box plot comparing position correlations of matched same-animal pairs (n=9) and unmatched cross-animal pairs (n=18). Mann-Whitney U test: U=162.0, p=3.47e-05***. Single data points overlaid on top. **d,** PCA on cropped and rotated mantle and head masks: average of highest and lowest 50 PC scores and difference map along each component for PC1 (top) and PC2 (bottom). PC1 distinguishes animals swimming horizontally (high) versus away or towards the camera (low), whereas PC2 distinguishes a more compact (high) versus streamlined body configuration (low), often seen during high speed swimming bouts (jet propulsion). **e,** Top: Speed (black) and PC2 (blue) time series; bottom: Speed versus PC2 (Pearson’s r=-0.574, p < 0.001). **f,** Experimental arena with two *T. helgolandica* annelids. **g,** Head-tail extraction from OCTRON-predicted masks: left, skeletonized centerline (white), endpoints classified as head (magenta) and tail (cyan) using local mask thickness; right, centerline (white), endpoints, reference point for head angle (black) and head direction (arrow) overlaid on raw frame. Confidence values ([0,1]) indicate certainty of head-tail assignment. **h,** Open-field tracks per annelid (same data as in f and g): line width indicates speed, hue indicates curvature direction and its strength; background is an average frame. **i,** Detrended head angle (black) and speed (pink) over time for one example animal. **j,** Left: Head-angle power spectral density (PSD, peak ≈1.9 Hz) for animal in (i). Right: box plot of peak frequencies across annelids (n=5 animals); median (pink line), interquartile range (gray box), and whiskers extending to min/max values to disable outlier classification. Individual data points were overlaid as dots.

While pixel-accurate masks are essential for analyzing body shape and appearance, many behavioral experiments - particularly long recordings, or crowded arenas containing many individuals - only require tracking positions and identities over time. For these applications, OCTRON includes a dedicated object-detection mode that converts SAM-generated masks directly into bounding-box training data, eliminating segmentation overhead at inference time to enable rapid, near-real-time tracking across large numbers of individuals. Across two model sizes (YOLO11l and YOLO11m) and two hardware platforms (Apple M2 Max/MPS and NVIDIA RTX PRO 6000 Blackwell/CUDA), detection-only models processed videos up to 23% faster than their segmentation counterparts, while maintaining comparable precision (mAP@50: 99.4% vs. 99.4% for YOLO11l, and 99.3% vs. 99.0% for YOLO11m, detection vs. segmentation respectively). The detection mode is therefore ideal for fast, multi-object tracking when information about shape is not required. For more complex scenarios, such as crowded scenes or multiple species, more powerful models are needed. We implemented two RT-DETR versions (RT-DETR-l and RT-DETR-x) that outperform YOLO in both speed and accuracy^9^. We validated multi-species detection by training a single RT-DETR model on SAM segmentation data of multiple individuals of two species (*Stentor coeruleus* and *Daphnia pulex*) swimming in the same arena (Fig. S3a-c). On held-out test frames, the model detected both species with high accuracy (mAP@50: 96.7% overall; 98.3% for *S. coeruleus*, 95.1% for *D. pulex*; precision 97.2%, recall 97.0%), confirming reliable simultaneous multi-species detection. However, multi-animal tracking introduces additional challenges: when an individual is briefly occluded, trackers may split its track into multiple fragments. To address this, we built a napari widget, allowing users to visually inspect and join candidate track fragments based on spatiotemporal proximity, improving track continuity (Fig. S3e-h, Video S3). Together, these features, fast detection-only models and interactive track curation, enable reliable detection and consistent tracking of multiple species with a single unified pipeline.

Not all animals have body features that make their orientation or posture easy to infer. Some, such as ctenophores (Fig. 1c), lack a distinct “anterior-posterior” axis, while in certain annelids (Fig. 2f), the head is so small relative to the body that achieving the image resolution needed for reliable detection is impractical. However, the segmentation masks produced by OCTRON enable keypoint-like extraction of distinct body points during secondary analysis. For example, after skeletonizing YOLO-predicted masks of a pelagic annelid (*Tomopteris helgolandica*), the centerline endpoints can be classified as ‘head’ or ‘tail’ based on the median medial-axis thickness of each endpoint (Fig. 2g), since the head region is rounder and bulkier than the tail. Similarly, access to the full shape of the animal allows assigning head and tail points with a confidence score derived from the observed head-to-tail asymmetry (Fig. 2g). The overall accuracy of these extracted head and tail points is similar to the accuracy achieved with an anchor point based model trained with DLC on the same frames used for OCTRON model training. However, since OCTRON point predictions are bound to the animal mask, whereas DLC predicts over the whole frame, fewer severe mispredictions occur with OCTRON (Fig. S4a,b). One strategy to limit erroneous tracking predictions is to filter out predictions below a confidence or likelihood cutoff (Fig. S4c,d). Because DLC keypoint likelihoods are often high even for off-animal false positives, eliminating severe jumps required strict filtering (>0.9 likelihood), which discarded a large fraction of frames and required extensive linear interpolation (Fig. S4c,d). In contrast, at YOLO’s standard 0.25 instance-confidence threshold, OCTRON retained nearly all frames while producing coherent trajectories with minimal interpolation (Fig. S4c,d), preserving the temporal continuity needed to resolve fine-scale swimming kinematics (Fig. 2h). Specifically, when swimming, *T. helgolandica* exhibits lateral body undulations^18^ that produce a rhythmic head-scanning motion (Fig. 2i, Video S4). By extracting points at fixed fractions along the centerline to define head angle relative to the anterior body axis, we quantified this oscillation across five animals, revealing a stable peak near 2 Hz (Fig. 2j). This is in line with known kinematics for this species^19^ and might have important implications for how *Tomopteris* process sensory stimuli during locomotion. Thus, in addition to analysis of full body changes, the rich shape information provided by OCTRON also enables secondary keypoints-like analysis of behavior.

### OCTRON enables tracking of camouflaging animals and extraction of body patterns

Finally, we evaluated OCTRON on an exceptionally challenging segmentation task: camouflaging *Sepia officinalis* cuttlefish. Individual animals were placed in an open, circular tank divided into three equally-sized areas that contained images of coarse, medium, and fine sand on its floor and walls. This arena type is designed to evoke distinct camouflage body patterns (disruptive, mottle, and uniform, respectively; Fig. 3a,b, S5a,b)^20–24^. SAM2 proved highly effective at segmenting mantle and head regions from minimal user input, dramatically reducing annotation time and enabling rapid generation of a large training dataset (Video S1). We found that our YOLO-trained model was able to reliably extract head and mantle regions across frames spanning a range of positions and skin patterns (Fig. 3b, S5b, and Video S5). By segmenting out the background (Fig. S5b,c), we were able to precisely align the tank regions and animal position data across videos (Fig. S5d). We next used the extracted head and mantle masks to derotate cuttlefish across all frames (Fig. 3c). A subset of those rotated and cropped cuttlefish frames were used to train a convolutional autoencoder (Fig. S6a) to extract 32 latent embeddings, which were reduced to two dimensions via principal component analysis (PCA) for visualization and clustering. Unsupervised clustering with HDBSCAN assigned cluster labels to all frames (Fig. S6b,c). These skin pattern clusters were then manually grouped into two classes: one combining predominantly mottle patterns with those extending into uniform patterns (merged due to the limited expression of purely uniform patterns in our experiment; Fig. S6c) and a second class capturing a variety of disruptive patterns (Fig. 3d, S6c). By combining pattern class information (Fig. 3d) with position data (Fig. 3e, top), we could precisely map the spatial occurrence of the two pattern classes and normalize by occupancy across the environment (Fig. 3e, bottom). As predicted, we found that mottle/uniform camouflage patterns were typically expressed in more uniformly patterned sectors of the arena (fine and medium sand), whereas a disruptive pattern appeared preferentially over the coarse background (Fig. 3f). OCTRON thus enables quantitative analysis of body patterns in addition to position, revealing behavioral insights inaccessible to location-only methods.

**Fig. 3:**
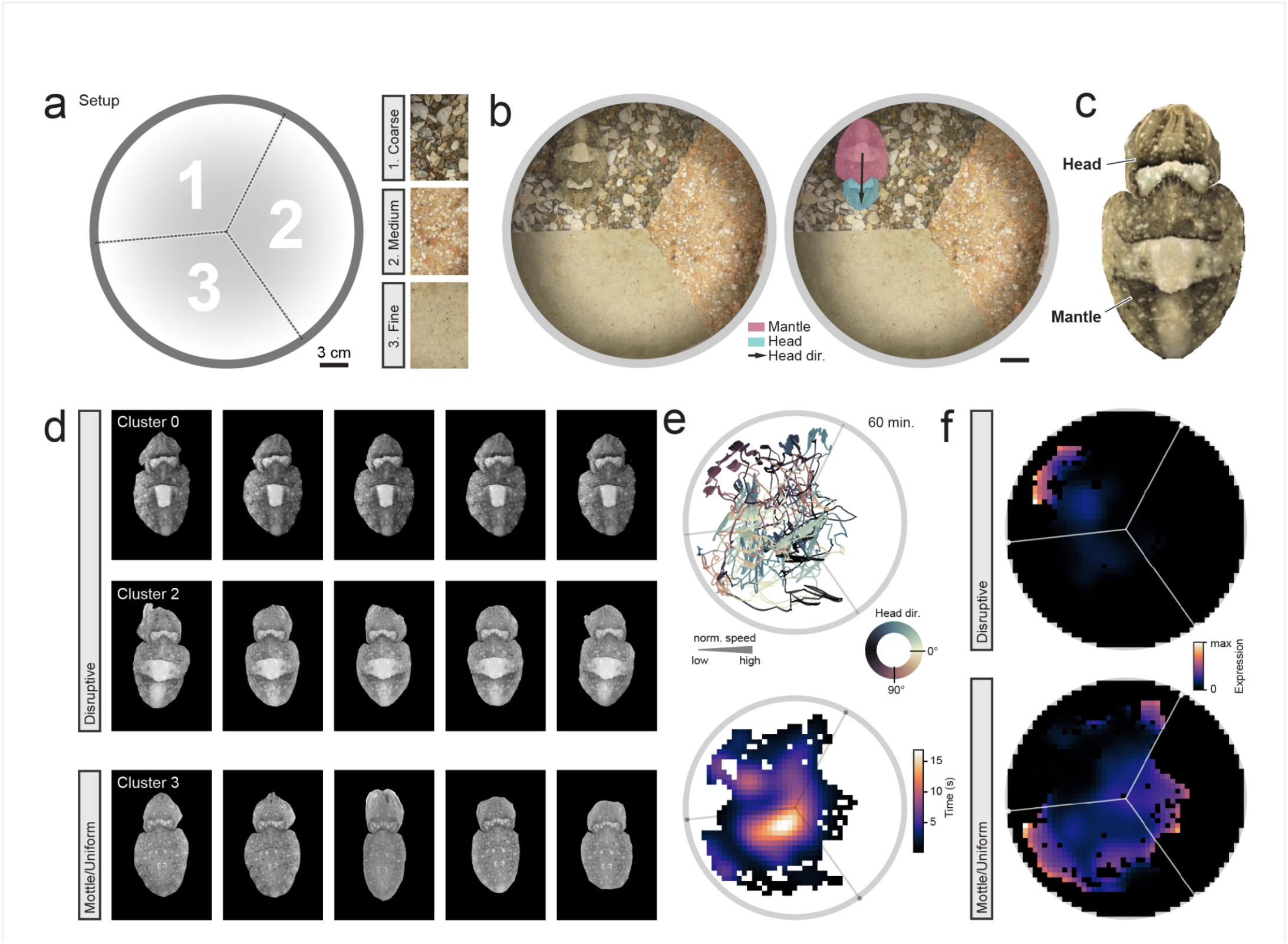
Tracking a camouflaging animal. **a,** Circular arena divided into three parts with images of various substrates (coarse, medium, and fine sand) covering the floor and walls. **b,** Example frame (left) and segmentation overlays (right) showing mantle (pink), head (cyan), and head direction (black arrow) of *Sepia officinalis*. Scale bar: 3 cm. **c,** Cropped and derotated dorsal view of cuttlefish from the image in (b). **d,** Example cropped and derotated L* channel frames from autoencoder clusters assigned to disruptive (clusters 0, 2) and mottle/uniform (cluster 3) pattern groups. **e,** Top: 60 min animal trajectory (pooled data from six individual videos). Color indicates head direction; line width indicates normalized speed. Bottom: heatmap of occupancy within the arena. **f,** Pattern tuning maps: disruptive (top) and mottle/uniform (bottom) expression probability normalized by occupancy; warmer colors indicate more frequent expression of the pattern.

## Discussion

By combining powerful, state-of-the-art tools for the rapid annotation, segmentation, detection, and multi-object tracking of animals, as well as visualization and export possibilities in a user-friendly graphical user interface or via the command-line, OCTRON provides a complete pipeline for markerless segmentation and tracking. The rich information supplied by the generated masks can benefit the behavioral analysis of all species, but are particularly well suited for animals that have complex, deformable body plans or blend into the background. While OCTRON is not the first pipeline to enable segmentation or tracking in soft-bodied animals^25–28^, it is the first fully trainable pipeline for multi-object segmentation tracking that is generally applicable across experimental setups and entities. For example, OCTRON may also be a valuable tool for analyzing timelapse videos of migrating cells: generated masks of (sub) cellular structures can be used to quantify positional and morphological changes, as well as to extract expression levels of fluorescent labels within.

While segmentation masks provide valuable information about body shape and can be used to generate keypoints-like skeletons, they make analyzing complex poses - such as joint position - challenging. Such analyses are still best achieved using keypoint-based methods. Importantly, the YOLO segmentation models that OCTRON uses already support keypoint tracking in principle, features we plan to integrate in the future. More broadly, computer vision is evolving rapidly, and specific model families will inevitably be superseded. OCTRON is built with longevity and maintainability in mind: because its annotation, training, and tracking stages are modular and independent from each other, future foundation models for annotation, object detectors, or multi-object trackers can replace or complement SAM, YOLO, RT-DETR, and BoxMOT without altering the user workflow or data formats. Together with its command-line interface and standardized outputs, this modularity also makes OCTRON ready for integration into automated, agentic analysis pipelines. Moreover, because OCTRON stores annotations in portable, hash-indexed Zarr archives that can be directly merged across users and laboratories, community adoption can rapidly build the cross-laboratory mask datasets needed to train pan-species segmentation "super-models" or domain-specific vision foundation models - analogous to recent pretrained pose-estimation efforts such as SuperAnimal^29^.

We also envision the pipeline supporting real-time on-device tracking through the less resource demanding detection models, enabling closed-loop experiments. Finally, we aim for OCTRON to support multi-camera angle data, enabling 3D reconstruction of body shapes and movements, and thereby bringing volumetric behavioral analysis within reach of animals with malleable bodies.

## Supporting information

Video S1 - Annotation workflow

Video S5 - Sepia officinalis

Video S4 - Tomopteris

Video S3 - Prediction cleaner widget

Video S2 - Octopus paralarvae

## Methods

### Pipeline

OCTRON consists of several modules, each of which harnesses powerful openly available tools. The implementation of these tools in each step of the OCTRON pipeline is described below.

#### Project organization

OCTRON operates on data contained within a user-defined project folder that consolidates all annotation and training data for a given project. The workflow begins with the user selecting a video for annotation, which can optionally be re-encoded with ffmpeg (https://www.ffmpeg.org) before being imported into OCTRON. During the import process, OCTRON creates a dedicated subfolder for the video inside the project directory, named according to a file hash of the video itself. This mechanism prevents mix-ups and safeguards data integrity, even if a newer video with the same filename unintentionally replaces an older one in the project folder. Annotation subfolders from multiple users who have annotated videos on their own machines can be incorporated into the project. This is achieved simply by copying and pasting the annotation folders along with their corresponding video files into the original project directory. This approach allows larger projects to be easily distributed across multiple users and later consolidated for the training stage (see below).

#### Annotation Module

OCTRON uses SAM models for semi-automated annotation of video frames. Users select from four model variants (SAM 2 Base Plus^5^, SAM 2 Large^5^, SAM 2 Large HQ^4^, SAM 3^6^) based on available GPU resources, with larger models and specialized high-quality (HQ) decoders providing greater precision. A fifth model, SAM 3 multi, enables rapid annotation of multiple objects in the same layer, facilitated by the use of concept prompts. Annotation is performed by one of two methods: point-based prompts, where users left-click to include regions and right-click to exclude them, and shape-based prompts, where users draw outlines (rectangles, circles, or polygons) around objects of interest. SAM generates region predictions (masks) in real-time based on user input, which are automatically saved to disk. Batch (forward) prediction functionality extends single-frame annotations across subsequent frames, with adjustable skip intervals to accelerate annotation of static or slowly-changing subjects. The annotation GUI organizes labels through a layer manager, where each label is associated with a point/shape layer for user input and a mask layer displaying SAM’s predictions. Annotations are stored as compressed zarr archives (python zarr ≥ 3.0, https://zarr.readthedocs.io/en/stable/), and label and video metadata are organized in JSON format. Users can reopen previously annotated videos and continue working on them. Also, SAM’s memory can be reset if batch predictions slow down due to accumulated frame context, while keeping annotations and mask layers intact.

#### Model Training Module

Training data generation splits annotated frames into training, validation, and held-out test subsets using a block-based, episode-aware strategy. Because SAM-assisted annotation (forward mask propagation) lets users label many frames quickly, annotated frames are often temporally correlated and clustered into distinct annotation bursts ("episodes") across a video; a naive per-frame random split would place near-duplicate neighboring frames on opposite sides of a split, inflating validation/test performance. OCTRON instead segments annotated frames into episodes at large temporal gaps, divides each episode into contiguous blocks (∼20 frames by default), and pools blocks across all episodes into a single train/validation/test budget (default 70%/15%/15%) so that realized split proportions match the requested fractions regardless of episode number or size. Validation and test blocks are then spread evenly across the recording timeline (stratified sampling) to remain representative, and a small buffer of frames (default: 1) is dropped at each boundary between differently-assigned blocks to introduce a temporal gap and further reduce leakage. This process is deterministic and reproducible via a fixed random seed. Optional pruning further excludes frames where not all labeled objects are present, and watershedding separates touching instances of the same object class. OCTRON trains two families of YOLO instance segmentation models that can also be used for detection, as well as an RT-DETR model family for detection only. The YOLO models, YOLO11^7^ and YOLO26^8^, each come in two size variants: medium (recommended for initial training) and large (for increased accuracy with sufficient training data). The RT-DETR model^9^ also comes in two size variants: large (recommended for real-time detection) and extra large (for even higher accuracy). Training parameters include image size (640, 1024, or a custom number of pixels divisible by eight, selected based on native video resolution and desired level of detail), number of epochs (with automatic early stopping after 100 epochs without improvement), and save period (checkpoint frequency). Training progress is monitored via terminal output or optionally through MLflow^30^ for real-time metric visualization. Evaluation metrics include mean Average Precision (mAP) at Intersection over Union (IoU) thresholds of 0.5 and 0.5:0.95, precision-recall curves, confusion matrices, and validation batch predictions. All training outputs, including model weights (epoch checkpoints [.pt], best.pt, and last.pt), per-epoch training metrics (results.csv), and diagnostic images (results.png, confusion_matrix.png, val_batchX_pred.jpg), are saved in the project folder’s model/training subdirectory.

#### Video Analysis Module

The analysis module applies a user-trained model to new videos using BoxMOT^31^ implemented tracking algorithms. Six tracking methods are available: motion-only trackers (ByteTrack^11^ and Observation-centric SORT^13^, computationally efficient but limited ability to track similar-looking objects) and motion+ReID trackers (Bot-SORT^12^, Deep OC-SORT^15^, Hybrid-SORT^14^, and BoostTrack++^16^, which incorporate visual feature matching for robust identity preservation during occlusions and interactions). Users can tune tracker parameters, enable single-subject mode (bypassing multi-object tracking), and activate detailed mode to enable the extraction of additional morphological features (via *skimage.measure.regionprops*^32^). Analysis parameters include morphological opening (disk radius for noise reduction), confidence threshold (0 to 1, filtering out low-confidence detections), intersection-over-union (IoU) threshold (controlling overlapping mask separation), and frame skip interval (for accelerated processing of videos that, for example, contain little changes over time). Output data is organized per video in an *octron_predictions* folder, with each analyzed video generating a subfolder containing per-track-ID CSV files and zarr-compressed mask archives. CSV files include frame indices, track IDs, label names, confidence scores, bounding box coordinates, and (in *detailed* mode) mask area, centroid position, eccentricity, solidity, and orientation derived from region properties. Prediction results can be visualized directly in napari with adjustable trajectory length and colormaps.

#### Track-curation interface

Multi-object trackers occasionally lose an individual for one or more frames - for instance during brief occlusions by conspecifics or arena features - and re-acquire it under a new identity, fragmenting a single animal’s trajectory into two or more tracks. Trackers may also produce spurious tracks from false detections. To resolve both issues, OCTRON provides a dedicated *Prediction cleaner*, implemented as a separate napari viewer dock widget that opens automatically when prediction results are loaded (or manually from the napari Plugins menu) (Fig. S3g,h; Video S3). Users select a source track from a dropdown-menu listing all tracks, colour-coded by label and annotated with the fraction of the video each track covers; low-coverage tracks, which are most likely fragments, appear at the top of the list. For a given selected source track, OCTRON automatically proposes join candidates: tracks sharing the same label that do not overlap the source in time and whose start or end points lie within a threshold distance of the source in both time and space (adjustable by the user). Candidates are listed together with the coverage gain that joining would yield, and the timeline can be jumped directly to the frame at which two tracks abut. Selecting a candidate triggers an interactive preview. The source and candidate tracks are isolated in the viewer and rendered with a colour gradient that converges on the putative join point, so that continuity in position and heading across the junction can be judged visually. Confirmed joins are written to disk as a new track: the .csv files of all member tracks are concatenated, re-sorted by frame index, and written under a new track ID; for segmentation projects, a corresponding mask array is created in the results Zarr store by copying all annotated frames from each member track. Tracks can also be discarded entirely by deleting the corresponding napari layer, which OCTRON detects automatically. All operations are non-destructive and fully reversible, with per-operation undo and the ability to perform a full-session reset.

#### File System Organization

The project folder contains annotation subfolders (one per video, named by abbreviated video file hash) with label-specific Zarr archives (*label_name masks.zarr*), video frame data (*video data.zarr*), and metadata (*object_organizer.json*, *video_info.txt*). Hash-based naming ensures video-annotation correspondence, preventing loading of video data that does not match the original annotation data exactly. Training generates a model subfolder containing *training_data* (train/test/val splits plus *yolo_config.yaml*), and a training subfolder populated during training with weights (*best.pt*, *last.pt*, epoch checkpoints), metrics (*results.csv*), and evaluation plots. Analysis outputs are stored in an *octron_predictions* folder within the video directory, structured as *video_file_tracker*/*label_track_id.csv* plus *predictions.zarr* and *predictions_metadata.json*. All data uses zarr ≥ 3.0 for compressed storage of masks and frame data, with CSV files providing tabular feature extraction compatible with pandas or other analysis libraries.

### Accuracy evaluation

To generate an independent ground truth dataset for model evaluation, random frames were extracted as follows; For every species-specific project (videos of *Ascarosepion bandense, Sepia officinalis, Mnemiopsis leidyi, Hydra vulgaris, Octopus vulgaris, Tomopteris helgolandica*), 25 frames were randomly sampled without replacement from each of several videos and loaded into a stack. Ground truth annotations were generated by manual annotation (labels layer, brush tool) of every frame in the stack in napari. Multiple annotators (n=5) independently painted binary masks for each label.

To assess model performance as a function of training data quantity, multiple YOLO11m models in segmentation mode were trained on subsets of the full training set. The complete training dataset was split into train/test/val partitions using fixed fractions (70%/15%/15%). From the training partition, five subset sizes were randomly sampled: n=60, 120, 240, 480, and 960 frames (each maintaining class balance). Models were trained for 350 epochs with maximum input image size of 640 pixels in one dimension, and cosine learning rate scheduling (initial lr=0.01). Mosaic augmentation was applied with probability 0.25 and disabled during the final 10 epochs to stabilize training. Data augmentation was applied within YOLO11 and comprised: (1) Color space jitter: HSV shifts with ranges ±0.25 for hue, saturation, and value channels; (2) Geometric transforms: random rotation ±180°, translation ±10% of image dimensions, scaling ±25%, shear ±2°, no perspective distortion, horizontal and vertical flips each with probability 0.5; (3) Advanced augmentations: mosaic (probability 0.25, combining 4 images), mixup (probability 0.25, alpha-blending two images), copy-paste (probability 0.25 in mixup mode, pasting objects from one image onto another). Masks were generated with overlap_mask=true to preserve overlapping instances and mask_ratio=2 for higher-resolution mask outputs during training. Model checkpoints were saved every 10 epochs, with best-performing models retained for evaluation. Model predictions were generated for the human-annotated test frames using each trained model. Frames were passed through YOLO segmentation with confidence threshold 0.5. For frames containing multiple detections of the same class, masks were combined via logical OR to produce a single unified mask per class, and confidence scores were averaged. Intersection over Union (IoU) was computed as the ratio of overlapping pixels to total pixels covered by either mask: IoU = |A ∩ B| / |A ∪ B|, where A is the predicted mask and B is the human-annotated mask. IoU was calculated frame-by-frame for each annotator separately. Additionally, an "average human annotation" was created by averaging binary masks across all annotators and thresholding at 0.5. Model predictions were compared against this consensus annotation to obtain IoU relative to collective human agreement. Frames lacking predictions for a given class were assigned IoU = NaN. IoU distributions were summarized as mean ± standard deviation across frames for each training set size and label class (Fig. 1f, S2). To compare model performance against single-annotator versus consensus annotations, independent-samples t-tests were used to assess significance (alpha=0.05).

### Animals

To demonstrate the versatility of OCTRON, we curated videos from our own work and one openly available dataset representing a variety of species, as described below in alphabetical order.

#### Ascarosepion bandense

Juvenile (3-4 months old) *Ascarosepion bandense* cuttlefish, also known as dwarf or stumpy-spined cuttlefish, were sourced from the Cephalopod Initiative breeding program at the Marine Biological Laboratory (MBL), Woods Hole, MA. In the absence of nationwide regulations for coleoid cephalopods, all procedures followed MBL’s Institutional Animal Care and Use Committee (IACUC) approved protocols under veterinary supervision. Stress was minimized throughout experiments, and animal health was monitored daily; any signs of stress (inking, excessive movement) paused experiments until recovery. Cuttlefish were housed individually in white, food-grade fiberglass pans (McMaster-Carr 4727T76-4727T761) divided into three compartments with opaque corrugated plastic sheets, within a closed-loop seawater system (“A Frame”) with a >30-gallon sump supplying all tanks. Salinity (31–33 ppt) was checked daily, and evaporative loss was compensated with filtered, desalinated water. The animals were fed twice daily with shrimp (*Palaemonetes spp.*). Behavior was recorded using overhead cameras (Basler acA2040-55uc, Thorlabs MVL25M1 lenses) mounted above enclosures at 20 fps (1544 × 1544 px). Frames and environmental data (temperature, brightness) were time-stamped via Bonsai (https://bonsai-rx.org; ^33^).

#### Hydra vulgaris

Dataset collected and published by Han et al.^34^ and openly available on Dryad (https://datadryad.org/dataset/doi:10.5061/dryad.f6v067r). The animals were imaged from above at 5 fps with a sCMOS camera attached to a dissecting microscope.

#### Mnemiopsis leidyi

Cydippid-stage *Mnemiopsis leidyi ^35^* from the F71 laboratory strain (a gift from Dr. William Browne, University of Miami) were reared at the University of California, Berkeley. Animals were maintained in artificial seawater (Instant Ocean dissolved in distilled water, adjusted to a salinity of 26 ± 2 ppt) and fed a diet of S-type rotifers and tiger pods (*Tigriopus californicus*) until they reached the lobate (adult) stage, approximately 1 cm in length. Once the animals were the desired size, they were placed in a tightly sealed 35 mm Petri dish connected to a syringe filled with artificial seawater. Videos were taken with a Sony HDR-CX900 camera at 720 x 720 pixel resolution at 30 fps for 7 min. Video contrast was enhanced using Adobe Premiere Pro.

#### Octopus vulgaris

Embryos of *Octopus vulgaris* were obtained from Eduardo Almansa (CSIC, Instituto Español de Oceanografía, Tenerife, Spain). Upon arrival in Belgium, they were maintained in a closed standalone system at the Laboratory of Developmental Neurobiology, KU Leuven, where subsequent experiments were conducted. All animals used in this study originated from a single spawning event in November 2022. Following hatching, paralarvae were kept unfed in beakers under low-light conditions. For behavioral recordings, one to three paralarvae (∼2 mm mantle length) were placed in a transparent 3 cm diameter Petri dish positioned on a custom-made adjustable transparent stage (Thorlabs). The stage was uniformly illuminated from below using a red light panel (ELF-200X200RD, Vision Light Tech). Recordings were captured from above using an enclosed monochrome CMOS camera with a trigger (PL-D734MU-NIR-T, Pixelink), equipped with a modular lens system (Navitar). The setup was enclosed by black curtains to eliminate ambient light interference. A red light filter was applied during acquisition, resulting in grayscale video recordings. Videos were recorded at a resolution of 2048 × 2048 pixels and a frame rate of 30.00 frames per second using Pixelink Capture software.

All procedures involving paralarvae were approved by the KU Leuven Ethical Committee for Animal Experimentation (permit P099/2021) and conducted in accordance with Directive 2010/63/EU.

### OCTRON analyses

We trained two YOLO segmentation models (YOLO11l at 1024 px and YOLO11m at 640 px) in OCTRON on 7,425 annotated images across 28 videos of swimming *Octopus vulgaris* paralarvae. For multi-animal tracking we used Hybrid-SORT in combination with “osnet_x1_0_market1501.pt” for Re-ID. For the trajectory plots in Fig. 2a we extracted spatial paths of identified mantle-head pairs (those with high correlation over the whole video) over a specified history window (500 frames, ∼16.7 seconds) ending 10 frames before the displayed frame. Background images were generated from single video frames with a circular mask applied to isolate the arena region. Trajectories were rendered using matplotlib^36^ *LineCollection* with line width proportional to mantle area (normalized to 99th percentile, scaled ranging 0.5-25 pixels) and color representing instantaneous speed. Line segment opacity increased linearly from 0 to 1 across the history window to emphasize recent movement. Position and area values were smoothed with Gaussian filters (σ=3 for position). For individual animal visualizations (Fig. 2a, right column), cropped regions were extracted centered on the midpoint between mantle and head positions. Frames were contrast-adjusted using 0th and 99.5th percentile values as vmin/vmax. Binary masks for mantle and head were extracted from segmentation outputs, converted to polygon coordinates using the imantics library (https://github.com/jsbroks/imantics), smoothed with a Gaussian filter (σ=2), and plotted as closed contours. Direction arrows indicating head-to-mantle orientation were drawn from mantle centroid to head centroid.

### Mantle-Head Pairing

To identify which mantle and head tracks belonged to the same individual, we computed Pearson’s correlations between all mantle-head position combinations separately for X and Y coordinates. Pairs exhibiting correlation >0.9 in both dimensions were classified as high-correlation pairs representing individual animals; all other combinations were retained as controls. Correlation matrices were visualized as heatmaps with high-correlation pairs positioned on the diagonal (Fig. 2b). Position correlation values (mean of X and Y correlations) were pooled across three separate datasets with video lengths 1:39 min, 5 min and 5 min (Fig. 2c).

### Morphology analyses

For each identified mantle-head pair, masks from both body parts were combined in each frame, rotated to a standardized "head-up" orientation using *scipy.ndimage.rotate* with nearest-neighbor interpolation and cropped to minimum bounding box. Up to 2500 frames per pair were processed. All rotated masks from all pairs across all datasets were flattened and combined into a single data matrix. PCA was performed using scikit-learn^37^. The fitted PCA transformation was then applied individually to each pair’s mask set to obtain PC scores for every frame. For visualization, frames with the highest and lowest 50 PC scores along each component were averaged and subtracted from each other (Fig. 2d). Then, the mantle position was differentiated and converted to speed (pixels/frame). PC2 scores were aligned to corresponding frames via frame index matching. Time series of speed and PC2 were smoothed with a Gaussian filter (σ=6) and plotted on dual y-axes for a representative pair (Fig. 2e, top). Pearson’s correlation between log₁₀-transformed speed and PC2 scores was computed across all frames from all pairs. A linear regression in log-speed space (y = PC2, x = log₁₀(speed)) was fitted to visualize the relationship (Fig. 2e, bottom). Scatter plots display every 15th data point for visual clarity.

### Comparison of detection and segmentation

To compare inference speed and precision between segmentation and detection, we trained the same two architectures on the identical annotated images in either segmentation or detection mode. Model accuracy is quantified with mAP@50 (mean average precision at an IoU threshold of 0.5), a single score summarizing both detection completeness (finding true objects) and localization accuracy (correctly placing their bounding boxes)^38^. Inference speed was benchmarked by running OCTRON’s full prediction and tracking pipeline (Hybrid-SORT; see below) on a held-out 1024×1024 video (300 frames, 3 repeats per model) on two hardware configurations: an Apple M2 Max (MPS backend) and an NVIDIA RTX PRO 6000 Blackwell (CUDA backend).

#### Sepia officinalis

Four juvenile *Sepia officinalis* were hatched from wild eggs collected in southern England and reared at the Marine Biological Laboratory Marine Resources Center (Woods Hole, MA, U.S.A.). At ∼9 months old, each animal was housed individually in an enriched environment with 15 °C perfused seawater, following MBL’s Cephalopod Care Policy. Trials were conducted in an isolated tank containing a cylinder (diameter: 27 cm) with tri-patterned laminated substrates (fine, medium, and coarse sand) on the walls and floor. Black fabric enclosed the setup to block external stimuli. An iPhone 14 mounted above the cylinder filmed each 10-minute trial at 1080p/30 fps under ring-light illumination.

### OCTRON analyses

We trained the YOLO “L” model (YOLO11l) in OCTRON in segmentation mode on 1158 images across a subset of 4 videos of *S. officinalis*. For analyses in Fig. 3, six videos of length 10 minutes of one animal were processed using OCTRON to identify the mantle and head regions. Detected regions were tracked across frames, and smoothed via Gaussian smoothing (σ = 2). For each frame, we extracted crops centered on the mantle region, rotated to align the mantle-to-head axis vertically, and registered them to a common canvas (600 × 400 pixels, Fig. 3c,d, S5b, S6c). For skin pattern analysis, we extracted the lightness (L\**) channel in CIELAB* color space, replaced background (outside-mask) NaN pixels with 50 (mid-range L*), and normalized values to [0, 1] by dividing by 100 (Fig. 3d, Fig. S6c). A symmetric convolutional autoencoder with a 32-dimensional bottleneck (convolutional layers, batch normalization, and ReLU activations) was trained on 1,000 randomly sampled frames (70% training, 30% validation/test) for 50 epochs using Adam (lr = 2×10⁻⁴, weight decay = 1×10⁻⁵) and mean squared error loss (Fig. S6a). We then passed the 1,000 frames through the trained encoder, projected the resulting 32-dimensional embeddings to 2D via principal component analysis (PCA), and fitted an HDBSCAN^39^ clusterer (min_cluster_size = 20, min_samples = 10, with prediction_data=True; label −1 treated as noise) on the 2D PCA coordinates (Fig. S6b). To classify skin patterns across all six recordings, cropped and masked frames (sampled every 10th frame) were normalized and passed in batches of 32 through the frozen encoder, projected into the fitted 2D PCA space, and assigned cluster labels via HDBSCAN approximate prediction (hdbscan.approximate_predict). We then manually associated cluster labels to two pattern classes (disruptive and a combined mottle/uniform class, reflecting the absence of a sharp distinction and the weak expression of fully uniform patterns in our experiment, Fig. S6c). For arena alignment across the six videos, video frames were analyzed to detect the circular tank boundary using ground segmentation masks from OCTRON predictions (Fig. S5c). For each video, we fitted a circle to the arena boundary by identifying three cardinal points (transitions between fine, medium and coarse background regions) and calculating the best-fit circle through these points. To enable cross-video comparisons, we then computed affine transformation matrices that aligned each arena to a reference frame (the first video in the dataset). The reference arena’s center and boundary points were used as the target, and transformations were calculated through affine transformations in *scikit* image^32^. An average circle was computed by transforming all individual arena circles to the reference frame and calculating the mean center and radius. Mantle and head coordinates from OCTRON predictions and centroid-based tracking were transformed to the reference frame using the pre-computed affine matrices (Fig. S5d). The center point between mantle and head was computed as the midpoint of the two body regions and used for all subsequent position analyses. Aligned tracking data and pattern classifications were merged on frame index. Occupancy maps were computed by binning animal positions into 45 bins (in x and y) within the circular arena and smoothed with Gaussian filtering (σ = 2 bins). Occupancy was normalized by frame time (1/30 s) to yield time spent in each spatial bin (Fig. 3e). Pattern-specific tuning maps were then generated by filtering tracking data by pattern class (disruptive or mottle/uniform) and computing occupancy maps restricted to frames where each pattern was expressed. Tuning maps were normalized by the overall occupancy map to yield the probability of expressing each pattern at each spatial location. Gaussian smoothing (σ = 2 bins) was applied to reduce noise in the final tuning maps (Fig. 3f).

#### *Stentor coeruleus* and *Daphnia pulex*

*Stentor coeruleus* and *Daphnia pulex* were selectively isolated from natural freshwater samples collected at Cape Cod, MA, under a stereomicroscope (AmScope SM-2T-LED). The isolated specimens were subsequently cultured and stably maintained in natural freshwater and spring water at 20-22 °C until the day of recording. For each behavioral recording, multiple specimens were placed in a custom, 3D-printed chamber (1.0 mm in diameter) with a cover glass glued to the bottom. A camera (Basler acA2040-55uc) was mounted on a stereomicroscope (Nikon SMZ18), and videos were acquired using Bonsai software^33^ at a frame rate of 20 Hz for 10 min at 1536 × 1536 pixel resolution. The field of view and image contrast were calibrated using Pylon Viewer.

### OCTRON analyses

Animals were automatically detected and tracked using OCTRON (v0.2.1). Annotations were generated from two recordings held out from the final behavioral dataset: masks for both species were seeded manually and propagated across frames using SAM2-assisted segmentation within OCTRON, then converted to bounding boxes for object-detection training. This yielded 609 annotated frames, split into training (n=410 frames), validation (n=99 frames), and held-out test (n=100 frames) sets, with two classes corresponding to *S. coeruleus* and *D. pulex*.

An RT-DETR-l object detector, initialized from COCO-pretrained weights, was fine-tuned at an input resolution of 1024×1024 pixels with a batch size of 8, an initial learning rate of 0.01 with cosine decay, and standard augmentation (mosaic, mixup, copy-paste, random rotation, flipping, and HSV jitter). Training was run on an NVIDIA GPU for up to 900 epochs and terminated after 210 epochs due to early stopping (100 epochs without validation improvement); the best checkpoint (epoch 110, selected by validation mAP@50-95) achieved a validation precision of 98.6%, recall of 97.9%, mAP@50 of 98.6%, and mAP@50-95 of 85.1%. To obtain an unbiased estimate of generalization performance, this checkpoint was then evaluated once on the fully held-out test set (n = 100 frames, 911 annotated instances, never used for training or checkpoint selection), achieving an overall precision of 97.2%, recall of 97.0%, mAP@50 of 96.7%, and mAP@50-95 of 84.2% (per-species mAP@50: 98.3% for *S. coeruleus*, 95.1% for *D. pulex*). This checkpoint was used for all subsequent analyses. The trained detector was applied to held-out recordings, and per-frame detections were linked into individual trajectories using the Hybrid-SORT multi-object tracker (BoxMOT) with an OSNet appearance re-identification model, run independently for each species. Automatically generated tracks were then manually reviewed in OCTRON’s track-curation interface, where fragments arising from brief tracking losses were merged into continuous per-individual trajectories.

#### Tomopteris helgolandica

Specimens were collected with plankton nets in the Trondheimfjord and the Stjørnfjord in Norway from 40-600 m depth. The specimens were housed in complete darkness in a temperature controlled room (12-14 °C) in fresh seawater pumped directly from the Trondheimfjord at 100 m depth. Once collected, the specimens were only handled under red and/or infrared light. A 3D-printed circular arena (Ø 105 mm, height 37 mm), designed in Blender 4.1 (Blender Online Community, 2024), was placed in an aquarium filled with freshly filtered seawater (5 µm polypropylene filter cartridge, 9-PA-5) covering the bottom 30 mm of the arena. To prevent the arena from floating, two external compartments were designed to hold small weights. The arena also accommodated LEDs inserted through 5 mm holes (spaced 7 mm apart and 20 mm above the bottom of the arena), controlled via an Arduino UNO microcontroller. The arena was illuminated from below with three infrared lights and filmed from above with an infrared camera (Basler acA2000-165umNIR with a 16 mm, 1:1.4 lens) at a resolution of 1000×1000 pixels with 16 ms exposure and a final frame rate of 7 fps.

### OCTRON analyses

We trained the YOLO “L” (YOLO11l) model in OCTRON in segmentation mode on 1660 images across a total of 14 videos of swimming *T. helgolandica*. For multi-animal tracking we used Hybrid-SORT in combination with “osnet_x1_0_market1501.pt” for Re-ID.

### Head-tail identification

Representative frames were analyzed to extract head and tail positions from binary animal masks. Masks were first skeletonized using morphological thinning to produce centerline path coordinates. The skeleton was converted to a pixel graph representation, and the longest path through the graph was identified using all-pairs shortest path algorithms. This longest path defined the animal’s primary body axis from one end to the other (Fig. 2g). Path coordinates were smoothed with a Gaussian filter (σ=3) to reduce pixelation artifacts. To discriminate head from tail, a distance transform was computed on the binary mask to obtain local thickness values at every pixel. For each endpoint of the longest skeleton path, a circular region (radius = 10% of mean major axis length across high-area frames) was defined. Within each circular region, the median distance transform value was calculated, representing the local body thickness. The endpoint with the greater median thickness was classified as the head (anterior end with larger diameter), while the opposite endpoint was classified as the tail. A confidence metric (*dist_conf*) was derived as 1 - (*thickness_ratio*), where *thickness_ratio* = *min(thickness_head, thickness_tail) / max(thickness_head, thickness_tail)*. To ensure reliability, detections with confidence below threshold (*dist_conf* < 0.35) were set to NaN and linearly interpolated with a strict limit of 10 consecutive frames to maintain temporal continuity without over-smoothing gaps. Optional Gaussian filtering reduced residual jitter in derived coordinates. An auxiliary angle reference point was defined by indexing into the reordered skeleton path at 20% of the total skeleton length from the head end (i.e., at position *−int(0.2 × skeleton_length)* in the tail-to-head ordered array). This reference point, located proximally along the anterior body, provided a stable local orientation vector (Fig. 2g). The allocentric angle was computed as arctan2(head_y-ref_y, head_x-ref_x), converted to degrees, and normalized to span the range [0°, 360°]. Body curvature was computed frame-by-frame by integrating perpendicular distances of skeleton points from the head-to-tail axis. For each skeleton coordinate, the cross product of the point vector (relative to tail) with the head-tail line vector determines whether the point lies to the left (positive cross product) or right (negative cross product) of the straight axis. The curvature metric was defined as (*left_ratio - right_ratio*), where *left_ratio/right_ratio* = fraction of skeleton points on each side. The metric also reported a straightness index computed as the ratio of Euclidean head to tail distance to total path arc length. Curvature time series were smoothed (Gaussian σ=3) for visualization (Fig. 2h). Composite backgrounds in Fig. 2h were created by randomly sampling 50 frames from the first 200 frames of each video, then averaging across time to produce an arena reference image. Animal paths were visualized using matplotlib^36^ *LineCollection*. Speed was computed from framewise centroid differences and Gaussian-smoothed (σ=2). Normalized speed scaled segment linewidths (range ≈1-19 pixels) to convey movement intensity. Head angle series (allocentric, arena coordinates) were unwrapped to eliminate 2π discontinuities, then detrended by subtracting a running median computed over a 5-second sliding window to remove slow drift while preserving oscillatory components. The detrended signal was plotted alongside Gaussian-smoothed speed (σ=2) on dual y-axes (angle: black; speed: pink) for a 20-second window (Fig. 2i). For frequency analyses, the instantaneous amplitude envelope and phase were computed via the Hilbert transform. Instantaneous frequency was derived as the time derivative of the unwrapped phase, scaled by fps and divided by 2π, then Gaussian-smoothed (σ=4). The amplitude envelope was similarly smoothed (σ=4) for stability. The power spectral density of the detrended head angle series was computed using Welch’s method with segment length *nperseg*=280 frames, overlap n=200 frames. Resulting power spectra were lightly smoothed (Gaussian σ=3) for visual clarity and plotted on linear frequency axes (1–3 Hz range) (Fig. 2j). Peak frequencies were identified using the FOOOF library^40^.

### OCTRON and DLC comparisons

Random frames (n=50 frames per video, 4 videos total) were extracted and used for (1) direct annotation of head and tail points in DLC (v.3.0.0) and subsequent training (pytorch backend, model: resnet50_gn, n=200 iterations with batch size 8 using standard DLC hyperparameters and augmentation [crop sampling 448 × 448 px, affine transforms, Gaussian noise σ = 12.75, motion blur]. The snapshot with the highest test mAP was retained [epoch 110; test mAP = 98.4%, RMSE = 2.43 px]), and (2) region annotations of the worm body in OCTRON and subsequent training (model: YOLO11l in segmentation mode, input image size: 1024), enabling the direct comparison of trained models in DLC and OCTRON. The 200 annotated frames were split 95/5 into training (n = 190) and test (n = 10) sets. The same extracted frames were also manually annotated in napari to establish ground truth head and tail positions. Three annotators independently marked head and tail point coordinates for each frame and these annotations were then averaged. Euclidean distances were calculated between OCTRON- and DLC-predicted and manually annotated head and tail positions. For head and tail point extraction in OCTRON the same method as described above (“Head-tail identification”) was used. For single frame comparisons in Fig. S4a,b, a baseline cutoff criterion of .25 confidence (OCTRON) or likelihood (DLC) was applied to filter out highly unreliable predictions, which retained 121 (out of originally 125) frames for comparison. For whole video comparisons (Fig. S4d), we analyzed one video (n=4479 frames) not included in the training dataset with both the OCTRON and the DLC model (using the best snapshot), and plotted the output over all frames. No static or dynamic cropping was applied; predictions were made on full 1000 × 1000 px frames. Data that did not meet confidence (OCTRON) or likelihood (DLC) thresholds (see filter criteria in Fig. S4d) were first replaced with NaN values and then linearly interpolated (violet points in Fig. S4d). Note that OCTRON confidence and DLC likelihood are defined on different scales: YOLO architectures train with task-aligned assignment^41^, producing a per-instance score that jointly reflects classification probability and predicted localization IoU (standard inference threshold: 0.25), whereas DLC outputs the maximum value of a per-keypoint spatial score map^1^ (default cutoff: 0.6, and commonly filtered at >0.9). Consequently, comparing fixed numerical cutoffs across Fig. S4c,d illustrates the trade-off between data retention and trajectory coherence across each method’s operating regime rather than a directly comparable probability threshold.

### Data processing and analyses

All analyses were performed in Python using numpy^42^, pandas^43^, scikit-learn^37^, scipy^44^, and matplotlib/seaborn^36,45^. Most plots used seaborn themes, and perceptually uniform colormaps via cmasher^46^.

### Statistics

Statistical tests were run using scipy^44^. When normality could not be assumed, non-parametric tests were used to provide more reliable results.

### Deployment options

OCTRON can be installed and run on macOS, Windows, and Linux operating systems.

## AI Use Disclosure

During the preparation of this manuscript and the accompanying analyses, the authors used Warp (Warp.dev, https://www.warp.dev), an AI-assisted development environment, to assist with editing portions of the manuscript text, developing benchmarking scripts, and executing model evaluation and speed-comparison analyses reported in this work. All AI-assisted output, including code, analyses, and text, was reviewed, verified, and edited by the authors, who take full responsibility for the content, accuracy, and integrity of the manuscript.

## Data availability

The A. *bandense* dataset was collected by Horst A. Obenhaus at the Marine Biological Laboratory in Woods Hole, MA, USA, and will be published elsewhere.

The *H. vulgaris* dataset has been published previously by Han et al ^34^ and is openly available on Dryad (https://datadryad.org/dataset/doi:10.5061/dryad.f6v067r).

The *M. leidyi* dataset was collected by Oscar M. Arenas at the University of California, Berkeley, and will be published elsewhere.

The *O. vulgaris* dataset was collected by Sofia Maccuro in the Laboratory for Developmental Neurobiology led by Eve Seuntjens, KU Leuven, Belgium, and will be published elsewhere.

The *S. coeruleus* and *D. pulex dataset* was collected by Takato Honda at the Marine Biological Laboratory in Woods Hole, MA, USA, and will be published elsewhere.

The *S. officinalis* dataset was collected by Johnston Renton at the Marine Biological Laboratory, Woods Hole, MA, USA, and will be published elsewhere.

The *T. helgolandica* dataset was collected by Nadia M. van Eekelen in the Marine Neuroscience Laboratory led by Ragnhild Irene Jacobsen at NTNU, Trondheim, Norway, and will be published elsewhere.

## Code availability

The OCTRON codebase and GUI is available on https://github.com/OCTRON-tracking and is documented on https://octron-tracking.github.io/OCTRON-docs.

## Acknowledgements

We thank May-Britt and Edvard Moser for supporting our pursuit of novel research directions, as well as colleagues at both the Kavli Institute for Systems Neuroscience and the Department of Biology at NTNU for invaluable technical support. In particular Kyrre Haugen, Klaus Jøran Jenssen, Endre Kråkvik, Haagen Waade, Dag Altin, Stig Koteng, and Siv Anina Etter. We also thank Carsten Wolff at the MBL for technical support, as well as Abhishek Kumar and Carsten Wolff for providing access to computing resources at the MBL. We thank Roger Hanlon at the MBL for his expert guidance on experiments that were conducted with *Sepia officinalis* in his lab and for providing data for this paper. We thank Lisa Abbo, David Remsen, Dan Calzarette, Nolan Gibbons, Brian McGonagle, and Jennifer Camerlengo from the Marine Resources Center at MBL for their support in setup building, maintenance and animal care, as well as Taylor Sakmar and Bret Grasse from the Cephalopod Initiative at MBL for their expert guidance in cephalopod care. We also thank Anne Sylvester and Briana Bertochi at MBL for their administrative support during the Grass and Whitman Fellowships (HAO, TH, OMA). We thank Eduardo Almansa for providing the *Octopus vulgaris* embryos and David Rowland for feedback on early versions of the manuscript. This work was supported by funding from Gidske and Peter Jacob Sørensens Fund (RIJ), Nansenfondet / Norsk Hydro (RIJ), Picower Fellowship from MIT and Freedom Together Foundation (TH), the Helen Hay Whitney Foundation (OMA), the Grass Foundation (HAO, TH, OMA), the Kavli Foundation (HAO, TH) and the MBL Whitman Centre (HAO, TH). SM and EvR were funded through KU Leuven grant C14/21/065 to ES and William Schafer and Fonds Wetenschappelijk Onderzoek grant G040124N to ES.

## Contributions

HAO designed and wrote the OCTRON pipeline, performed analyses and generated figures. Testing and feedback on early versions of OCTRON were done by RIJ, NMvE and LH. Experiments were designed and/or implemented by RIJ, NMvE, JRe, OMA, EAL, TH, SMa, and HAO, with supervision from RIJ, EAL, KB, ES, and HAO. Ground truth annotations were performed by RIJ, NMvE, LH, EvR, and JRi. OCTRON’s command line interface functionality was coded by MA. SMi performed codebase cleaning and implemented the testing suite. Writing was carried out by RIJ, NMvE, EvR, JRe, TH, ES, and HAO with feedback from all authors. Funding was acquired by RIJ, TH, OMA, ES and HAO.

## Supplementary figures

**Fig. S1:**
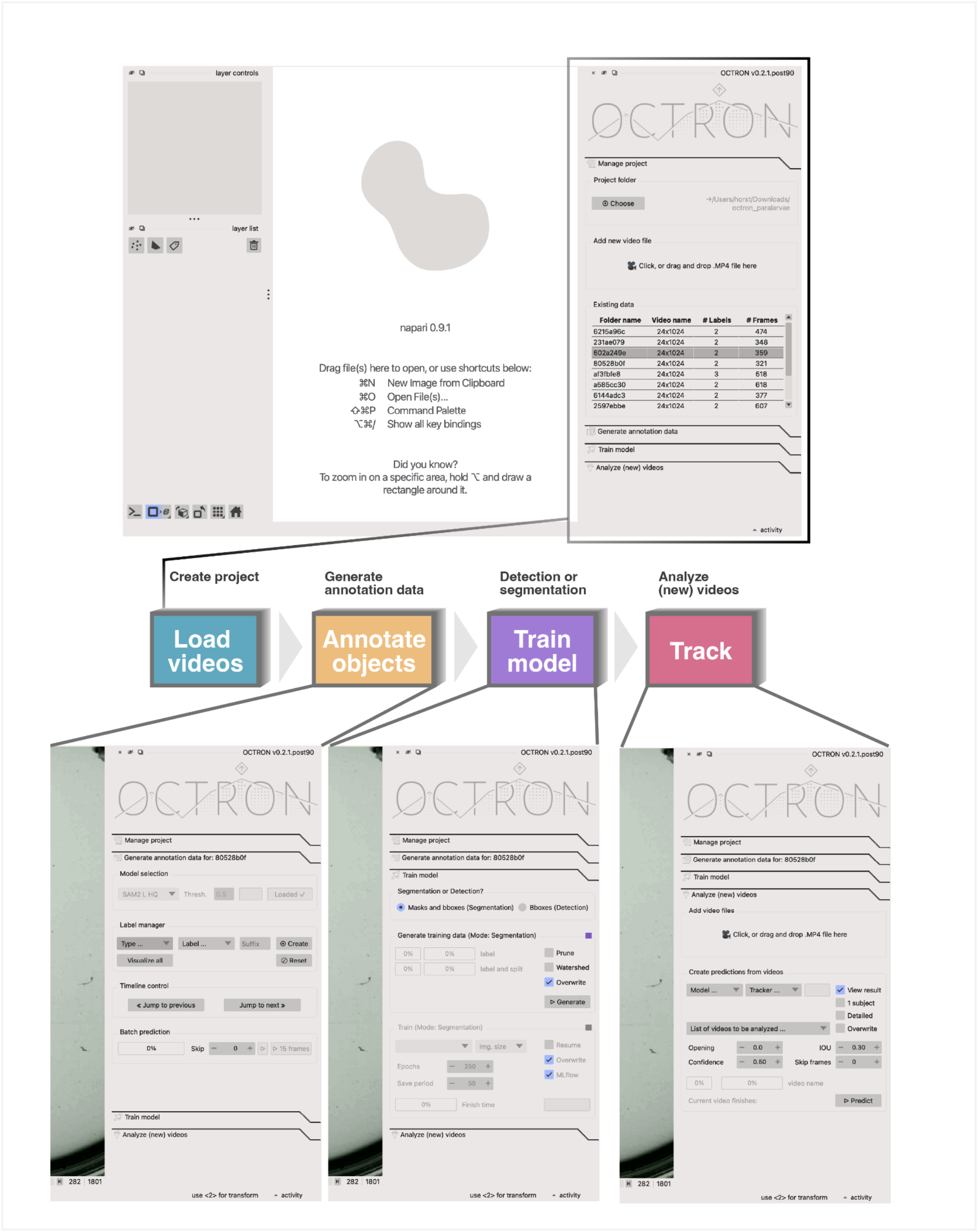
The OCTRON graphical user interface (GUI). Top: Screenshot of the OCTRON GUI after a project folder has been opened, showing the number of annotated frames for each video in the right-hand panel. Middle: Schematic illustrating the flow through the pipeline modules. Bottom: Screenshots of the right-hand panel in OCTRON, showing the options available in each module.

**Fig. S2:**
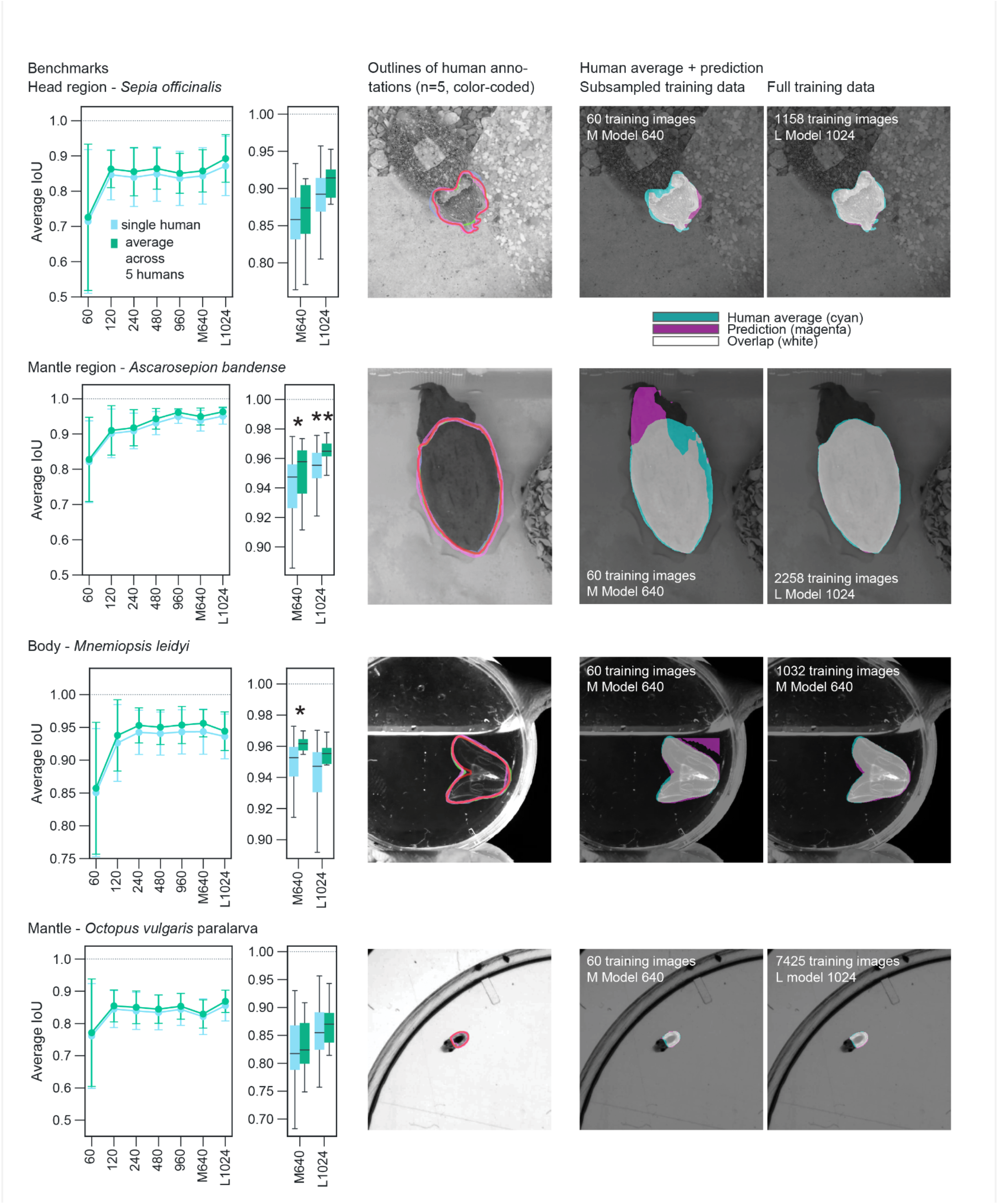
Additional benchmarking results across training sizes and models. As in Figure 1f. Segmentation accuracy versus human labels: Mean ± SD Intersection over Union (IoU) and box plots for models trained on varying sizes (YOLO M model trained on 60, 120, 240, 480, 960 frames) and on the full training dataset (YOLO M, and YOLO L at either 640px or 1024px max training image size) for four different species. Statistics: * p < 0.05, ** p < 0.01

**Fig. S3:**
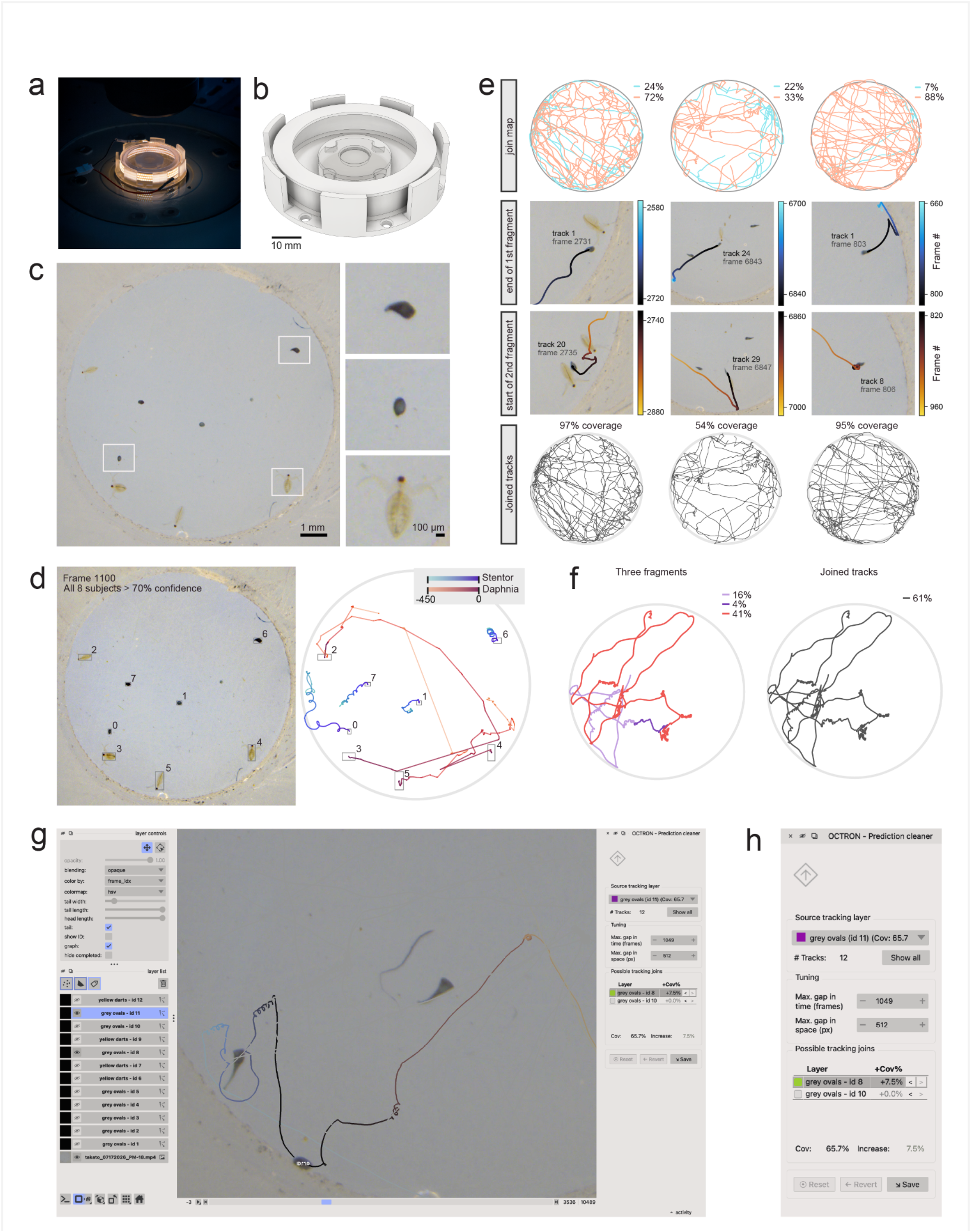
Recording setup for *Stentor coeruleus* and *Daphnia pulex,* and automated track curation pipeline. **a,** Recording chamber mounted on a stereomicroscope stage. **b,** 3D-printed chamber design (scale bar, 10 mm). **c,** Representative arena frame with *Stentor coeruleus* and *Daphnia pulex*; insets show individual specimens at higher magnification. **d,** Example frame with all eight tracked subjects exceeding 70% detection confidence (left) and their trajectories (right), colored by species and frame index (blue, *Stentor coeruleus*; orange, *Daphnia pulex*). Trailing traces are 450 frames (∼22s) long. **e,** Three representative track-joining examples for *S. coeruleus* recorded in three different sessions: raw fragment pairs before joining (first row), each track fragment labeled with its percentage of coverage in terms of number of frames ("join map"); zoomed in views of the join point — last frame of the earlier fragment (second row) and first frame of the later fragment (third row) — colored by frame number; and the resulting joined trajectory with its overall coverage in terms of number of frames (last row, "joined tracks"). **f,** Example of a three-fragment join ("3-partition") for one *S. coeruleus.* Left: the three raw fragments (colored, with individual coverage percentages as in e). Right: and the resulting single joined trajectory (grey, with overall coverage). **g,** OCTRON track-curation interface (built on napari) used to inspect and manually curate track fragments. Left: layer list with one layer per identified object (descriptive labels were used, where ‘dark ovals’ represent *S. coeruleus* and ‘yellow darts’ represent *D. pulex*), alongside standard (napari built-in) track-layer display controls (color, tail length/width, ID and overlays). Center: video frame with overlaid raw track fragments. Right: the fragment-joiner widget for the currently selected source fragment. **h,** Magnified view of the fragment-joiner widget showing the current source fragment and its frame coverage ("Cov"), the total number of raw fragments for that identity, adjustable maximum temporal and spatial gap thresholds used to search for candidate track fragments to join, a ranked list of candidate fragments with their predicted coverage gain (+Cov%), the resulting total coverage and increase, and reset/revert/save controls. Every step is reversible.

**Fig. S4:**
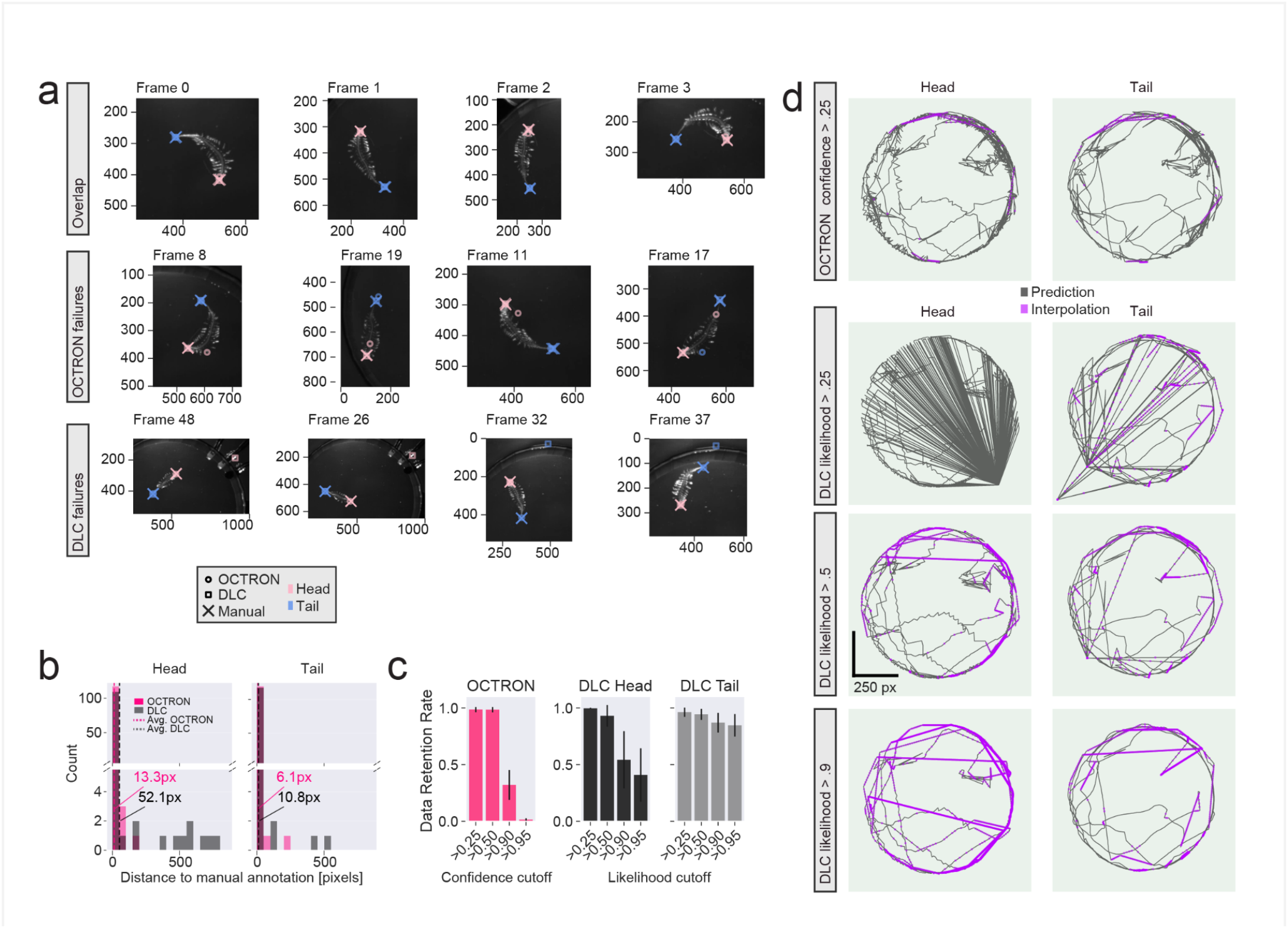
Comparing OCTRON and DeepLabCut (DLC) tracking performance. **a,** Representative examples showing overlays of OCTRON (confidence > .25) and DLC (likelihood > .25) predictions, and manual annotations. Top row: clean cases in which all three annotation types agree. Middle row: OCTRON failure cases. Note that erroneous predicted anchor points are still localized near the body of the animal. Bottom row: DLC failure cases. Note that erroneous predictions can appear anywhere in the arena, often far from the animal’s body. **b,** Distribution of distances between OCTRON (confidence > .25) or DLC (likelihood > .25) predicted anchor points and ground truth anchor points (average of three human annotators) across 121 labeled images. Stippled lines with associated annotations indicate population means. Bin size: 50 pixels. **c,** Proportion of retained frames across confidence (OCTRON, left) and likelihood (DLC, middle and right) cutoffs. Although both metrics span [0, 1], they are computed and calibrated differently: OCTRON reports YOLO’s per-instance confidence, a composite score reflecting both class probability and localization overlap (IoU) that is bounded below 1.0 for deformable objects and uses a standard deployment cutoff of 0.25. In contrast, DLC reports the per-keypoint maximum of a spatial score map, which is skewed toward high values even on false positives (see panel d at >0.25 and >0.5) and typically requires strict filtering (>0.6–0.9), at the cost of substantial data loss. **d,** Example predicted trajectories across an entire video (n=4479 frames) at different confidence/likelihood thresholds, with missing data interpolated (violet).

**Fig. S5:**
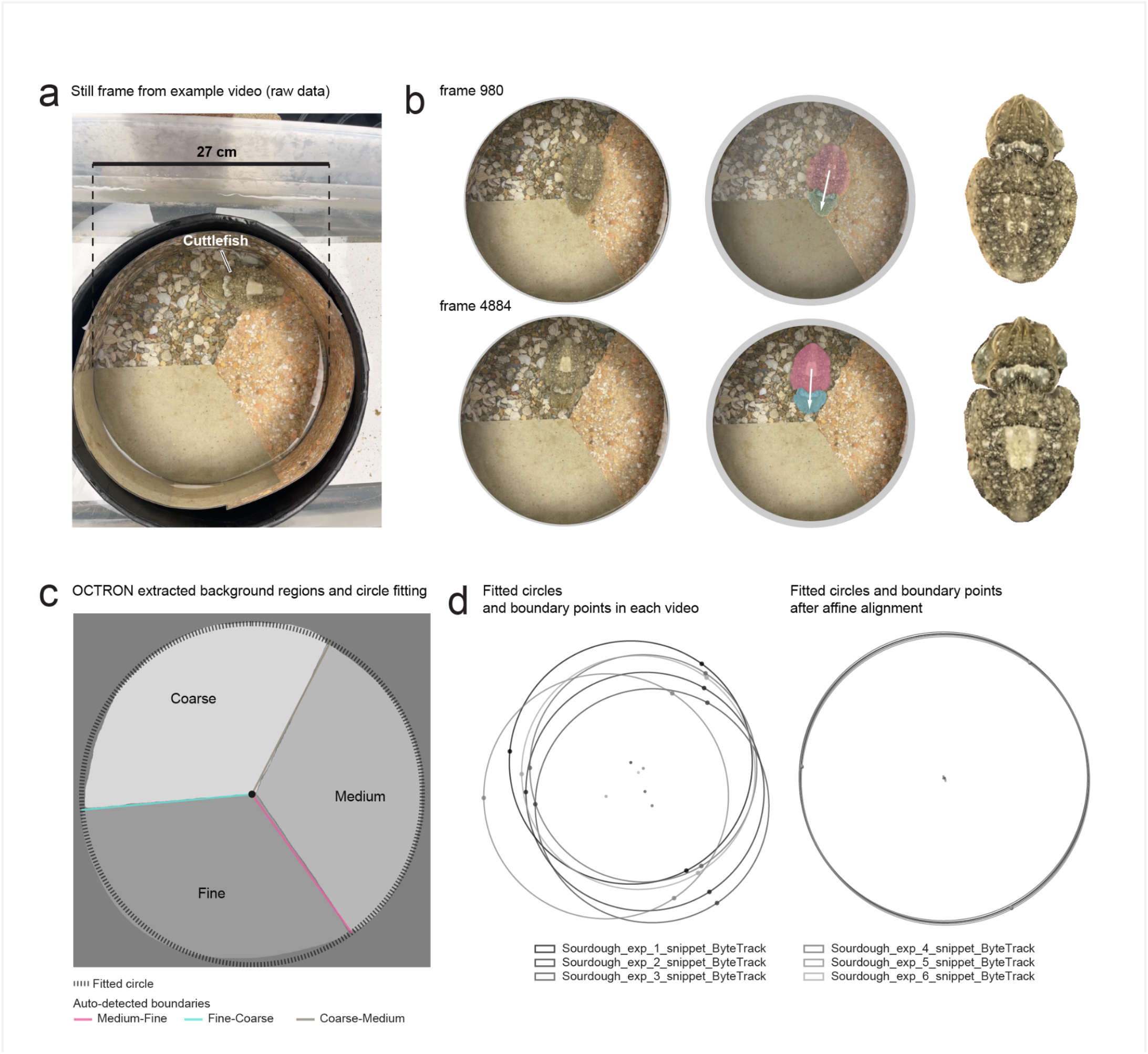
Example segmentations and arena alignment. **a,** Raw video frame of the circular arena with three substrate sectors and one *Sepia officinalis* cuttlefish. **b,** Example frames: raw view (left), OCTRON detections overlaid (middle; mantle in pink, head in cyan, white arrow = head direction), and corresponding cropped and derotated animal images (right). Top (video frame 980) middle image also shows the detected background regions in gray as an overlay. **c,** OCTRON-derived background masks with fitted tank boundary (dashed circle) and sector boundaries (Medium-Fine, Fine-Coarse, Coarse-Medium). **d,** Fitted tank boundaries and substrate boundary points for each video (left) and after affine alignment to a common reference frame (right); gray tones denote individual recordings.

**Fig. S6:**
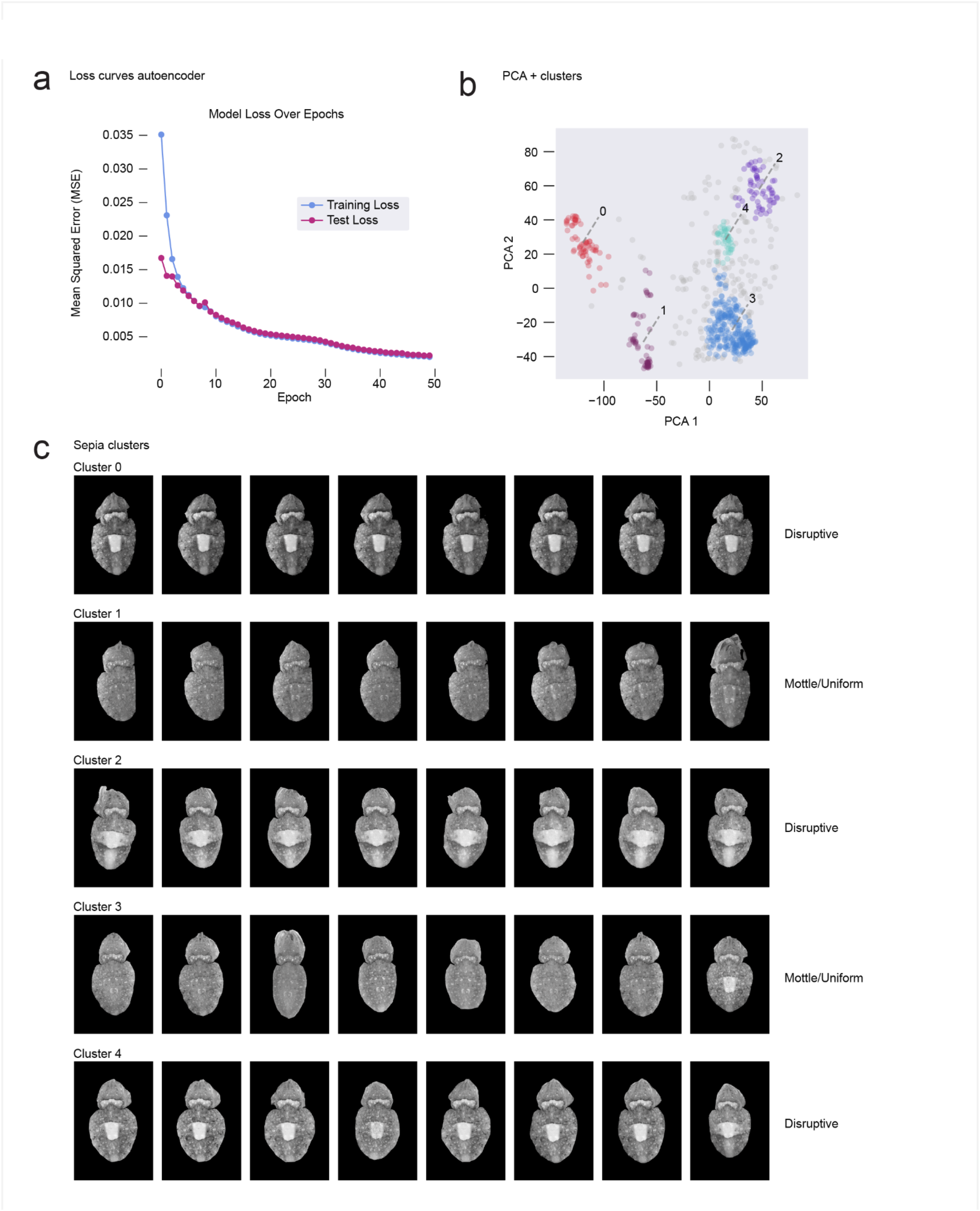
Autoencoder, clustering of patterns and clustered pattern examples. **a,** Autoencoder training and test loss (mean squared error, MSE) over 50 epochs. **b,** PCA projection of 32D latent embeddings with HDBSCAN cluster labels. **c,** Representative derotated L* channel frames from each cluster (0–4), illustrating disruptive (clusters 0, 2, 4) and mottle/uniform (clusters 1, 3) pattern groups.

## Supplementary videos

**Video S1 Demonstration of the annotation workflow in OCTRON**.

Demonstration of rapid annotation-data generation for a *Sepia officinalis* cuttlefish, from model and label creation, through manual single-frame annotation to automatic batch prediction across 15 frames.

**Video S2 Tracking three *Octopus vulgaris* paralarvae**.

Same data as shown in Fig. 2a. Mantle (pink) and head masks (green) are shown as overlays for each tracked octopus. Trailing paths are color-coded by octopus ID with saturation reflecting passed time. The data were tracked via HybridSort (see methods).

**Video S3. Prediction cleaner widget in OCTRON.** Demonstration of joining fragmented tracks: a source track is selected, candidate fragments close in time and space are inspected, and appropriate matches are joined to it.

**Video S4 Tracking two marine annelids (*Tomopteris helgolandica)***.

Same data as shown in Fig. 2f. Masks, IDs and trailing paths of mask centroids are shown for both annelids. Video is playing back at original speed (7fps). The extracted centreline (white), head (pink dot), tail (blue dot) and anchor point for head angle extraction (black dot) are shown as overlays for each animal.

**Video S5 Tracking a camouflaging cuttlefish (*Sepia officinalis*)**.

A single *Sepia officinalis* cuttlefish. Excerpt of data shown in Fig. 3. The masks of the head (green) and mantle (pink) regions are shown, along with the trailing paths of their respective centroids. Throughout the video, the cuttlefish undergoes multiple changes in its skin pattern.

## References

1. Mathis, A. et al. DeepLabCut: markerless pose estimation of user-defined body parts with deep learning. Nat. Neurosci. 1 (2018) doi:10.1038/s41593-018-0209-y.

2. Pereira, T. D. et al. SLEAP: A deep learning system for multi-animal pose tracking. Nat. Methods 19, 486–495 (2022).

3. Biderman, D. et al. Lightning Pose: improved animal pose estimation via semi-supervised learning, Bayesian ensembling and cloud-native open-source tools. Nat. Methods 21, 1316–1328 (2024).

4. Ke, L., et al. Segment anything in high quality. arXiv [cs.CV] (2023) doi:10.48550/arXiv.2306.01567.

5. Ravi, N., et al. SAM 2: Segment Anything in Images and Videos. arXiv [cs.CV] (2024) doi:10.48550/arXiv.2408.00714.

6. Carion, N., et al. SAM 3: Segment Anything with Concepts. arXiv [cs.CV] (2025) doi:10.48550/arXiv.2511.16719.

7. Jocher, G. & Qiu, J. Ultralytics YOLO11. (2024).

8. Jocher, G., et al. Ultralytics YOLO26: Unified real-time end-to-end vision models. arXiv [cs.CV] (2026) doi:10.48550/arXiv.2606.03748.

9. Zhao, Y., et al. DETRs beat YOLOs on real-time object detection. arXiv [cs.CV] (2023) doi:10.48550/arXiv.2304.08069.

10. Broström, M. & Kozak, M. BoxMOT: Pluggable State-of-the-Art Multi-Object Tracking Modules. (Zenodo, 2023). doi:10.5281/zenodo.8132989.

11. Zhang, Y. et al. ByteTrack: Multi-object tracking by associating every detection box. arXiv [cs.CV] (2021) doi:10.48550/arXiv.2110.06864.

12. Aharon, N., Orfaig, R. & Bobrovsky, B.-Z. BoT-SORT: Robust associations multi-pedestrian tracking. arXiv [cs.CV] (2022) doi:10.48550/arXiv.2206.14651.

13. Cao, J., Pang, J., Weng, X., Khirodkar, R. & Kitani, K. Observation-centric SORT: Rethinking SORT for robust multi-object tracking. arXiv [cs.CV] (2022) doi:10.48550/arXiv.2203.14360.

14. Yang, M., et al. Hybrid-SORT: Weak cues matter for Online multi-Object Tracking. arXiv [cs.CV] (2023) doi:10.48550/arXiv.2308.00783.

15. Maggiolino, G., Ahmad, A., Cao, J. & Kitani, K. Deep OC-SORT: Multi-pedestrian tracking by adaptive re-identification. arXiv [cs.CV] (2023) doi:10.48550/arXiv.2302.11813.

16. Stanojević, V. & Todorović, B. BoostTrack++: using tracklet information to detect more objects in multiple object tracking. arXiv [cs.CV] (2025) doi:10.48550/arXiv.2408.13003.

17. Sofroniew, N., et al. napari: A Multi-Dimensional Image Viewer for Python. (Zenodo, 2026). doi:10.5281/zenodo.3555620.

18. Gouveneaux, A., Gielen, M.-C. & Mallefet, J. Behavioural responses of the yellow emitting annelid Tomopteris helgolandica to photic stimuli. Luminescence 33, 511–520 (2018).

19. Daniels, J., Aoki, N., Havassy, J., Katija, K. & Osborn, K. J. Metachronal swimming with flexible legs: A kinematics analysis of the midwater polychaete Tomopteris. Integr. Comp. Biol. 61, 1658–1673 (2021).

20. Barbosa, A. et al. Cuttlefish camouflage: the effects of substrate contrast and size in evoking uniform, mottle or disruptive body patterns. Vision Res. 48, 1242–1253 (2008).

21. Allen, J. J., et al. Cuttlefish dynamic camouflage: responses to substrate choice and integration of multiple visual cues. Proc. Biol. Sci. 277, 1031–1039 (2010).

22. Chiao, C.-C., Kelman, E. J. & Hanlon, R. T. Disruptive body patterning of cuttlefish (Sepia officinalis) requires visual information regarding edges and contrast of objects in natural substrate backgrounds. Biol. Bull. 208, 7–11 (2005).

23. Hanlon, R. T. et al. Cephalopod dynamic camouflage: bridging the continuum between background matching and disruptive coloration. Philos. Trans. R. Soc. Lond. B Biol. Sci. 364, 429–437 (2009).

24. Chiao, C.-C., Chubb, C. & Hanlon, R. T. A review of visual perception mechanisms that regulate rapid adaptive camouflage in cuttlefish. J. Comp. Physiol. A Neuroethol. Sens. Neural Behav. Physiol. 201, 933–945 (2015).

25. Romero-Ferrero, F., Bergomi, M. G., Hinz, R. C., Heras, F. J. H. & de Polavieja, G. G. Idtracker.Ai: Tracking all individuals in small or large collectives of unmarked animals. Nat. Methods 16, 179–182 (2019).

26. Allan, D. B., Caswell, T., Keim, N. C., van der Wel, C. M. & Verweij, R. W. Soft-Matter/Trackpy: V0.7. (Zenodo, 2025). doi:10.5281/ZENODO.16089574.

27. Hebert, L., Ahamed, T., Costa, A. C., O’Shaughnessy, L. & Stephens, G. J. WormPose: Image synthesis and convolutional networks for pose estimation in C. elegans. PLoS Comput. Biol. 17, e1008914 (2021).

28. Banerjee, S. C., Khan, K. A. & Sharma, R. Deep-worm-tracker: Deep learning methods for accurate detection and tracking for behavioral studies in C. elegans. Appl. Anim. Behav. Sci. 266, 106024 (2023).

29. Ye, S. et al. SuperAnimal pretrained pose estimation models for behavioral analysis. Nat. Commun. 15, 5165 (2024).

30. The MLflow Development Team. MLflow. (2026).

31. Broström, M. BOXMOT.

32. van der Walt, S. et al. scikit-image: image processing in Python. PeerJ 2, e453 (2014).

33. Lopes, G. et al. Bonsai: an event-based framework for processing and controlling data streams. Front. Neuroinform. 9, 7 (2015).

34. Han, S., Taralova, E., Dupre, C. & Yuste, R. Comprehensive machine learning analysis of Hydra behavior reveals a stable basal behavioral repertoire. Elife 7, (2018).

35. Agassiz. On the metamorphoses undergone by certain Fishes before acquiring the adult form. Ann. Mag. Nat. Hist. 16, 69–70 (1865).

36. Hunter, J. D. Matplotlib: A 2D graphics environment. Computing in Science & Engineering 9, 90–95 (2007).

37. Pedregosa, F. et al. Scikit-learn: Machine Learning in Python. J. Mach. Learn. Res. 12, 2825–2830 (2011).

38. Everingham, M., Van Gool, L., Williams, C. K. I., Winn, J. & Zisserman, A. The pascal visual object classes (VOC) challenge. Int. J. Comput. Vis. 88, 303–338 (2010).

39. McInnes, L., Healy, J. & Astels, S. hdbscan: Hierarchical density based clustering. J. Open Source Softw. 2, 205 (2017).

40. Donoghue, T. et al. Parameterizing neural power spectra into periodic and aperiodic components. Nat. Neurosci. 23, 1655–1665 (2020).

41. Feng, C., Zhong, Y., Gao, Y., Scott, M. R. & Huang, W. TOOD: Task-aligned one-stage Object Detection. arXiv [cs.CV] (2021) doi:10.48550/arXiv.2108.07755.

42. Harris, C. R. et al. Array programming with NumPy. Nature 585, 357–362 (2020).

43. Team, T. P. D. Pandas-Dev/Pandas: Pandas. (Zenodo, 2020). doi:10.5281/zenodo.3509134.

44. Virtanen, P. et al. SciPy 1.0: fundamental algorithms for scientific computing in Python. Nat. Methods 17, 261–272 (2020).

45. Waskom, M. L. seaborn: statistical data visualization. Journal of Open Source Software 6, 3021 (2021).

46. van der Velden, E. CMasher: Scientific colormaps for making accessible, informative and “cmashing” plots. J. Open Source Softw. 5, 2004 (2020).

